# A minimally morphologically destructive approach for DNA retrieval and whole genome shotgun sequencing of pinned historic Dipteran vector species

**DOI:** 10.1101/2021.06.28.450148

**Authors:** Petra Korlević, Erica McAlister, Matthew Mayho, Alex Makunin, Paul Flicek, Mara K. N. Lawniczak

## Abstract

Museum collections contain enormous quantities of insect specimens collected over the past century, covering a period of increased and varied insecticide usage. These historic collections are therefore incredibly valuable as genomic snapshots of organisms before, during, and after exposure to novel selective pressures. However, these samples come with their own challenges compared to present-day collections, as they are fragile and retrievable DNA is low yield and fragmented. In this paper we tested several DNA extraction procedures across pinned historic Diptera specimens from four disease vector genera: *Anopheles, Aedes, Culex* and *Glossina*. We identify an approach that minimizes morphological damage while maximizing DNA retrieval for Illumina library preparation and sequencing that can accommodate the fragmented and low yield nature of historic DNA. We identify several key points in retrieving sufficient DNA while keeping morphological damage to a minimum: an initial rehydration step, a short incubation without agitation in a low salt Proteinase K buffer, and critical point drying of samples post-extraction to prevent tissue collapse caused by air drying. The suggested method presented here provides a solid foundation for exploring the genomes and morphology of historic Diptera collections.

**Significance statement:** Large museum collections of pinned insects could provide important snapshots of genomes through time, but unfortunately DNA retrieval from such fragile samples often leads to severe morphological damage, especially in delicate species such as disease transmitting Diptera. In this study we have worked on a combined method that minimizes morphological damage while maximizing the retrieval of DNA from dry pinned Diptera species. We identified the importance of tissue rehydration, gentle DNA lysis buffer incubation, and critical point drying to restore collapsed tissues. We hope this approach will make it possible for more historic insect specimens to become available for genomic research while ensuring they remain intact for morphological studies.

## Introduction

Over the past 100 years heavy use of pesticides has resulted in novel evolutionary pressures on targeted species, often leading to successful control that is swiftly followed by resistance (Forgash 1984; AL-Ahmadi 2019). For example, vector control measures such as insecticide treated bednets and indoor residual spraying have had a positive impact on the control of human malaria vectors of the *Anopheles* genus (Kleinschmidt & Rowland 2019). However, this also caused an increase in insecticide resistance (Forgash 1984). Present-day populations of major malaria vector species from the *An. gambiae* complex across sub-Saharan Africa now have widespread insecticide resistance (Kerah-Hinzoumbé et al. 2008; Edi et al. 2014; Clarkson et al. 2021; Munywoki et al. 2021), with similar increases observed in other malaria vectors, such as *An. funestus* (Riveron et al. 2015), *An. stephensi* (Yared et al. 2020), as well as other Dipteran disease vectors such as *Aedes aegypti* (Satoto et al. 2019), the vector responsible for transmitting yellow and dengue fever (Powell et al. 2018). Widespread insecticide use in sub-Saharan Africa started in the 1950s with DDT (dichlorodiphenyltrichloroethane) (Mendis et al. 2009), with new insecticides frequently introduced (Oxborough et al. 2015). While mosquito genetic population structure change has been researched over the past 20 years (Githeko et al. 1996; Gloria-Soria et al. 2016; Anopheles gambiae 1000 Genomes Consortium et al. 2017), it is unclear what population structure looked like prior to major vector control initiatives. Museum and other historic collections contain specimens pre- and post- the introduction of DDT and other insecticides, providing snapshots of populations that were reacting to these new evolutionary pressures in “real time”. These collections could be used to fill gaps in our understanding about the evolution of insecticide resistance and compare how historic populations compare to present-day genomic landscapes. Furthermore, historic collections also include the name-bearing type specimens for these species and recovering genomic data from them could help us understand complex species evolution (Strutzenberger et al. 2012; Prosser et al. 2016).

Museum Diptera specimens are often identified prior to pinning, and due to improper mounting, storage, and general wear and tear from handling can accumulate morphological damage (Walker et al. 1999). On top of the samples themselves being quite fragile, the DNA from historic insect specimens is more fragmented and damaged than DNA from present-day individuals of the same species, and yields are usually lower. After the death of an organism DNA start degrading due to chemical processes such as hydrolysis and oxidation, which cause strands to break and accumulate base damage, the most common being cytosine deamination into uracil, particularly in single stranded overhangs at the end of molecules (Lindahl 1993). DNA in ancient samples such as fossil bones and teeth accumulated thousands of years of postmortem damage and the fraction of 5’ C>T and 3’ G>A DNA substitutions resulting from cytosine deamination can reach 20%-60% depending on age and species (Briggs et al. 2007; Dabney et al. 2013). Because of that, specific techniques have been developed for the retrieval, sequencing and processing of ancient DNA, including recovery of ultrashort DNA fragments during extraction (Dabney et al. 2013; Rohland et al. 2018), optimization of double or single stranded library preparation and sequencing (Meyer & Kircher 2010; Briggs & Heyn 2012; Gansauge et al. 2020), as well as post-sequencing approaches to deal with high proportions of contaminant DNA and to make the most out of short, deaminated endogenous reads (Skoglund et al. 2014; Racimo et al. 2016; Link et al. 2017). Fortunately, historic samples had a shorter time frame for accumulating damage, are often stored in archives with stable temperatures and humidity levels limiting microbial growth and contaminant DNA accumulation, and while the retrieved endogenous DNA still tends to be very short, substitutions arising from cytosine deamination are much lower (about 2-5%) (Bi et al. 2013; Weiß et al. 2016; Gutaker et al. 2017; Parejo et al. 2020), making it easier to account for them during data processing such as variant calling. Previous work on historic insect specimens have primarily focused either on complete destruction of individual specimens or parts of specimens followed by PCR or whole genome sequencing (Parmakelis et al. 2008; Staats et al. 2013; Timmermans et al. 2016; Andrade Justi et al. 2021), or less destructive approaches and PCR based methods (Gilbert et al. 2007; Santos et al. 2018), which can lead to very high failure rates due to the fragmented nature of older DNA, as well as amplification of contaminant DNA molecules that are much longer than the target endogenous DNA. Some papers combine a minimally destructive approach with whole genome sequencing, but these were done on more robust insect species that can withstand harsher DNA lysis buffers (Tin et al. 2014; Parejo et al. 2020; Andrade Justi et al. 2021).

In this paper we present a minimally morphologically destructive approach for DNA retrieval from pinned historic vector Diptera specimens with the goal of maximizing DNA retrieval while minimizing irreversible morphological damage to precious specimens. Specimens were selected from the London Natural History Museum (NHM) Diptera collection. We focused primarily on sub-Saharan African malaria transmitting *Anopheles* mosquitoes and confirmed the range and efficacy on *Aedes, Culex* and *Glossina*. We couple this with ancient DNA purification techniques, library preparation optimized for low yield extracts with short inserts, and processing the sequencing data using ancient DNA pipelines. A schematic of the initial steps, from selecting, cataloguing, extracting DNA, purification and returning the specimens to the collection is summarized in Supplementary Figure S1. We show that by using this approach it is possible to retrieve nuclear data and consensus mitochondrial genomes with shallow shotgun sequencing.

## Results

### DNA retrieval from *Anopheles gambiae* complex mosquitoes within the last century

As there has been limited genomic work on historic *Anopheles* specimens (Parmakelis et al. 2008; Andrade Justi et al. 2021), we wanted to evaluate yields and ancient DNA characteristics in mosquitoes collected within the past century, as well as the stability of their morphological integrity during handling. For this we selected several major and minor vector species from the *An. gambiae* complex spread across six decades (1930s-80s) (specimen metadata in Supplementary Table S1). This initial approach included rehydrating pinned samples prior to submerging them in “lysis buffer A”, adapted from a recently published low salt Proteinase K tissue clarifying buffer for preparing samples for microscopy (Santos et al. 2018). After overnight incubation the specimens were rinsed with ethanol and air dried, while the lysis buffer was purified using a modified MinElute silica column approach used in ancient DNA research (Dabney et al. 2013). We noticed the rehydration step was crucial in order to minimize damage caused by static electricity, as samples after rehydration only occasionally lost legs or rarely the head, which was primarily due to the original placement of the specimen pin. We also noticed early on that air drying post extraction was not suitable, as very fine structures such as abdomens, limbs and antennae collapsed. To counteract this, the samples were taken through a series of ethanol concentrations from 30% to 100%, and then critical point dried (CPD) with liquid CO_2_. This procedure, although laborious, greatly improved morphological characteristic accessibility (Supplementary Figure S2).

In terms of DNA retrieval we had a wide range of estimated DNA yields (determined using a Quant-iT™ PicoGreen™ dsDNA Assay Kit), from only 3 ng for a specimen collected in the 1930s (value similar to our extraction blanks) to nearly 200 ng from 1980s specimens (Figure 1A, Supplementary Table S3), with much longer DNA fragments also present in the more recently collected specimens as assessed by an Agilent Bioanalyzer High Sensitivity DNA Analysis Chip (Supplementary Figure S3). These are estimates of the total DNA retrieved from each specimen, and are affected by co-extracted non-DNA molecules, a mixture of double and single stranded fragments, and could be microbial growth post-pinning.

**Fig. 1.**
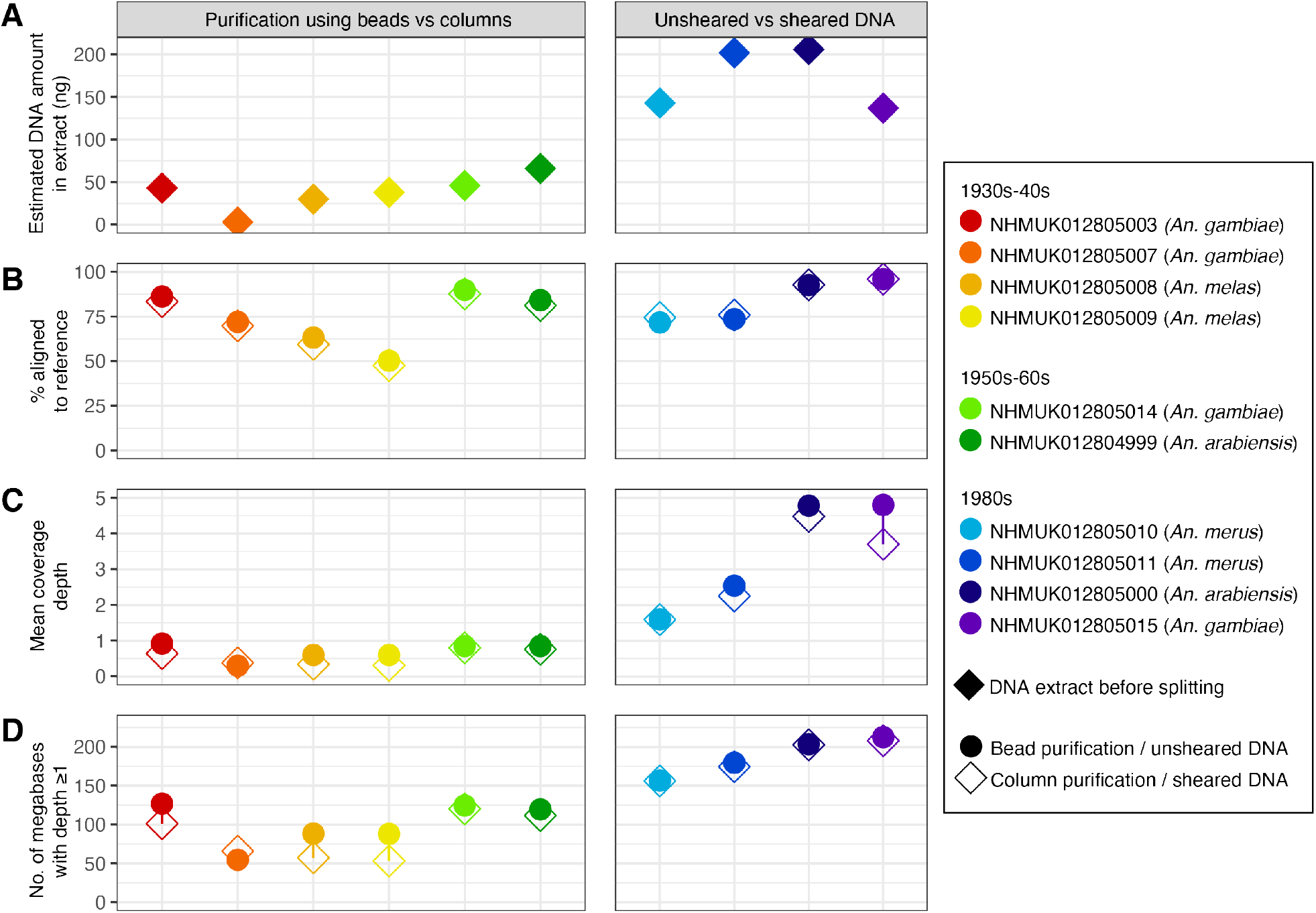
Sequencing data summary of 10 historic *Anopheles gambiae* complex specimens. For older samples (left and middle panels, 1930s-1960s) libraries post-ligation were split in half and purified using SPRI beads or MinElute silica columns, while for younger samples (right panels, 1980s) DNA extracts were split in half, one half was sheared, and the unsheared and sheared aliquots underwent library preparation separately. A) Estimated DNA yields in nanograms (ng) measured by Quant-iT™ PicoGreen™ dsDNA Assay Kit. B) Percentage of sequences in each library aligning to the AgamP4 *An. gambiae* reference. Variation can be driven by non-endogenous DNA and/or by aligning to a mismatched reference genome. C) Mean nuclear depth of coverage achieved for each library. D) Number of megabases in the AgamP4 reference genome covered with a depth of at least 1x or more (maximum 230,466,657).

Due to the fragmented and low yield nature of these DNA extracts we had to adapt our double stranded library approach to fit this type of input compared to present-day samples. Initial extract volumes were split into two library preparation strategies for each sample. For older samples that contained DNA below 300 bp based on Agilent chips the post-ligation libraries were purified either with SPRI beads or silica columns. For younger samples that contained DNA up to 10,000 bp the extract was split in half with one half being sheared and the other going into library preparation unsheared. Libraries were amplified by indexing PCR which introduced 8 bp tags on both ends, pooled and sequenced using a 75 paired-end (PE) approach on a lane of an Illumina HiSeq 2500 system (pool containing 20 sample libraries with a lower fraction for 4 extraction blank libraries). Libraries were then processed using the ancient DNA pipeline EAGER (Fellows Yates et al. 2021) as well as tools already integrated in samtools (Li et al. 2009).

The summary statistics after mapping to the *An. gambiae* reference genome (AgamP4) are shown in Figure 1 panels B-D. Several specimens investigated here are from species in the *An. gambiae* complex that are substantially diverged from the *An. gambiae* s.s. reference. Therefore, lower percentages of aligning reads could be driven by this divergence and/or non-endogenous DNA such as DNA from microbial growth. In this initial assessment we see that the percentage of DNA aligning to the reference varies in samples of more distantly related species (*An. melas, An. merus*), likely due to reduced similarity to the reference instead of an increased level of microbial contamination (Fontaine et al. 2015). For species more closely related to the AgamP4 reference (*An. gambiae, An. arabiensis*), we see a much higher fraction of sequences aligning, with older samples (collected further in time) showing slightly lower nuclear coverage compared to younger samples of the same species. We also see that the different approaches during library preparation are virtually indistinguishable (circles vs diamonds in Figure 1), so for library preparation throughout the rest of this study we opted for the most parsimonious protocol of no shearing along with bead purification prior to indexing PCR in order to minimize the number of steps and simplify multichannel or robot work by avoiding individual sample tubes.

As we expect our library DNA inserts to be short, we merged overlapping 75 bp paired-end reads (with a minimum 11 bp overlap), and retained both merged (inserts ≤139 bp) and unmerged paired reads (inserts >139 bp). When looking at the size distribution of aligned reads we notice that the fraction of 75 bp reads increases with age, going all the way up to 50-60% of total reads for samples from the 1980s, while the oldest samples contain inserts that are on average only 40-60 bp long (Supplementary Figure S4A, Supplementary Table S3). Through this set we also clearly illustrated the need to use the correct polymerase to PCR amplify the libraries post-ligation, as our 5’ C>T signals were affected by a polymerase that could not recognise uracils, the incidence of which was estimated to about 3-5% at molecule ends and 1-2% within the DNA fragments (Supplementary Figure S4B, C). The opposite strand’s 3’ G>A signal is not as affected since the pairing of adenines to uracils is done during library preparation.

### Maximizing DNA retrieval while minimizing morphological damage

Our second experiment focused on the retrieval of DNA and the level of morphological damage using three different lysis buffers (extensively tested on present-day *Anopheles* mosquitoes and outlined in (Makunin et al. 2021)) on *An. gambiae, An. melas* and *An. funestus* samples at the oldest age range of historic collections we are aware of (1920s-40s) (specimen metadata in Supplementary Table S1). Samples were initially incubated overnight, and we observed substantial levels of clarification and pigment loss, especially in smaller species such as *An. funestus* (Supplementary Figure S12), so a smaller subset of *An. funestus* specimens were instead incubated for only 2 h. Unfortunately, we could not perform a thorough assessment of morphology as the CPD instrument malfunctioned, and there was substantial damage caused by prolonged exposure to high concentrations of ethanol as well as insufficient CO_2_ during the drying procedure. However, it appeared that a 2 h incubation caused less damage and internal tissue loss (including blood meals) compared to an overnight incubation (Supplementary Figure S13).

Comparable to what was observed in present-day samples (Makunin et al. 2021), there are no clear differences in total DNA yields between the three buffers (Figure 2A, Supplementary Figure S6, Supplementary Table S3). It is far more likely that the observed variation in DNA retrieval is due to within-sample set variation, and not caused by the buffers themselves, though we have tried minimizing such variation by selecting samples collected from similar times and locations. Lysis buffer C and G containing no Proteinase K resulted in the lowest yields but a similar level of morphological damage, so Proteinase K was used in all further testing of these two buffers.

**Fig. 2.**
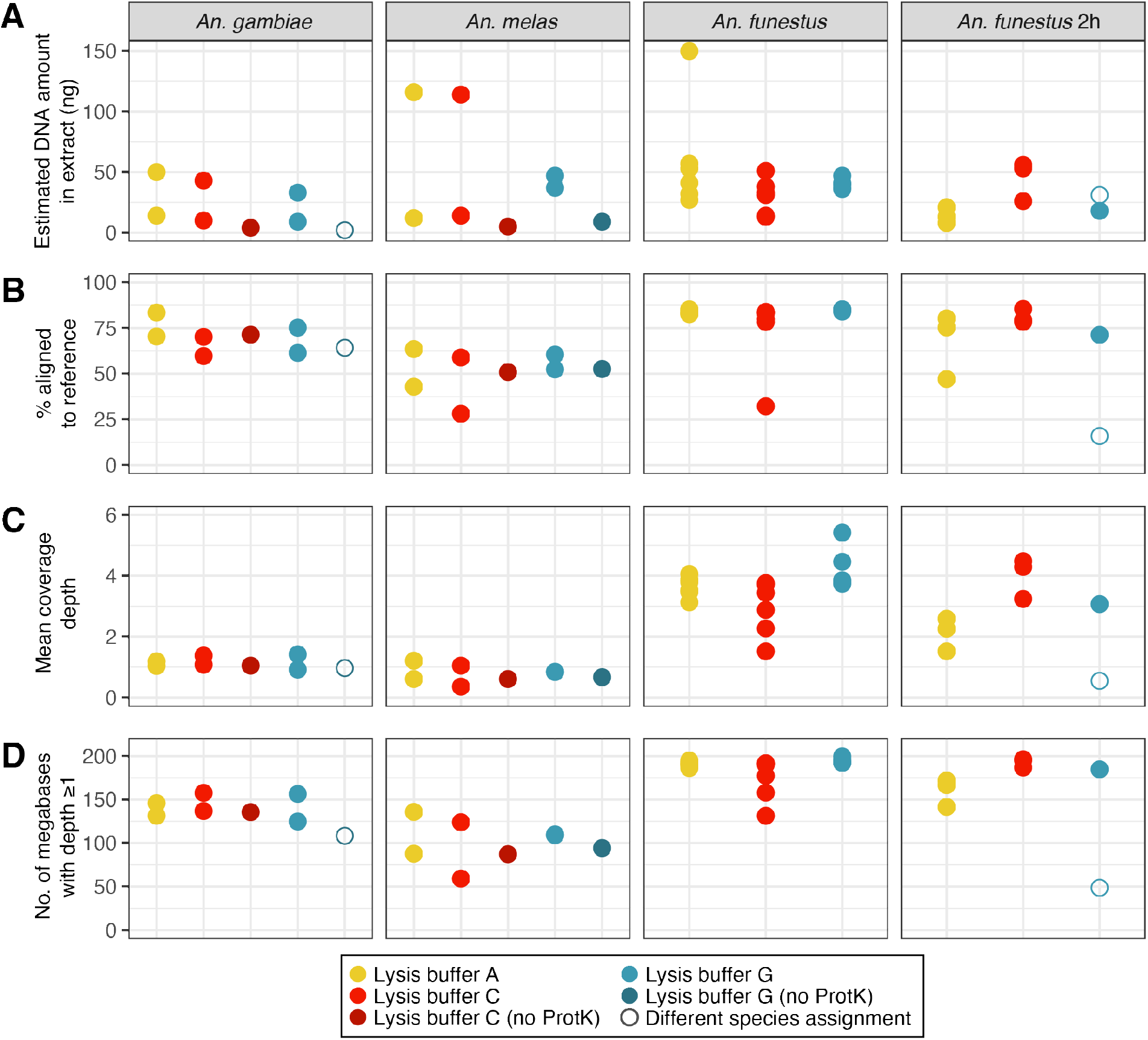
Sequencing data summary of overnight incubated *Anopheles gambiae* (n = 8), overnight incubated *An. melas* (n = 8), and overnight or 2 h incubated *An. funestus* (n = 26) specimens in three different lysis buffers, two of which were also tested when they contained no Proteinase K. A) Estimated DNA yields in nanograms (ng) measured by Quant-iT™ PicoGreen™ dsDNA Assay Kit. B) Percentage of sequences in each library aligning to *An. gambiae* (AgamP4) or *An. funestus* (AfunF3) reference. C) Mean nuclear coverage depth in each library. D) Number of megabases in the AgamP4 reference covered with a depth of at least 1x or more (maximum 230,466,657) for *An. gambiae* and *An. melas*, or AfunF3 (maximum 210,975,322) for *An. funestus*. Samples represented with empty circles were found to be a different species after sequencing and mitochondrial assembly (*An. funestus* mapped to AfunF3 in the first column, *An. rivulorum* in the last column).

Libraries were prepared following the simplified approach described above (no shearing, lower concentration SPRI purifications post-ligation and post-PCR), and samples were pooled into three pools (*An. gambiae/An. melas* pool 24 samples + 9 blanks, *An. funestus* overnight pool 18 samples + 3 blanks, *An. funestus* 2 h pool 20 samples + 3 blanks; not all libraries from the first and last pool are discussed here as they are part of another project). The pools were sequenced on three lanes of an Illumina HiSeq 4000 75 PE with the same processing and summary statistics generation as for our first experiment, *An. gambiae/An. melas* libraries were mapped to AgamP4, and the *An. funestus* libraries were mapped to the *An. funestus* nuclear (AfunF3) and mitochondrial (NC_038158.1) references.

After sequencing we noticed a similar picture to our first sample set, with *An. gambiae* samples on average mapping 70% and *An. melas* mapping 51% to the AgamP4 reference, while *An. funestus* samples on average had 79% mapping to the AfunF3 reference (Figure 2B, Supplementary Table S3). In this set we also found two samples, NHMUK010633485 previously determined as *An. gambiae*, and NHMUK013655440 previously determined as *An. funestus*, for which their original morphological assessment and consensus mitochondrial genome species were not a match. Mitochondrial data from these specimens grouped with mitochondrial genomes most similar to *An. funestus* and *An. rivulorum*, respectively (Supplementary Figure S9). The first sample was therefore mapped to AfunF3 and NC_038158.1, while unfortunately for *An. rivulorum* we do not have a more appropriate reference. Comparing the overnight and 2 h incubations for the *An. funestus* specimens, we retrieved a substantial amount of mosquito DNA after a 2 h incubation compared to overnight (similar total yields, percent aligned, and just slightly lower coverage) (Figure 2B-D), confirming a 2 h incubation is better for maximizing DNA retrieval from pinned specimens while minimizing morphological damage. We also see no substantial difference between buffers in terms of deamination patterns or retrieval of longer or shorter DNA sequences (Supplementary Figure S7). The difference in length observed for *An. funestus* samples, with plenty of inserts being too long to overlap and a prominent 75 bp peak, is likely due to the different strategy of SPRI bead purification post-ligation (one round of 2.2x SPRI compared to 2.5x SPRI used for the *An. gambiae* complex samples) and post-indexing PCR (two rounds of 1x SPRI compared to a single round of 1.2x SPRI) (Supplementary Table S3). This larger portion of longer molecules, combined with a lower pool plex likely explains why we obtain higher coverage for *An. funestus* samples compared to the *An. gambiae* complex samples (Figure 2C), and so a slight selection against very short molecules in samples of these decades and younger, coupled with a plex level of under 20 samples per pool, could be beneficial to lower sequencing costs.

### Efficiency across different vector Diptera species

In our final experiment we focused on what we believe is currently the best approach for simultaneously retrieving adequate amounts of DNA for further genomic work and minimizing morphological damage to the pinned dry specimen based on both our findings and current literature: rehydrating the pinned specimens prior to handling for 3 h at 37°C, incubating in lysis buffer for 2 h at 37°C, rinsing the specimens in 30% ethanol and storing in 50% ethanol prior to full ethanol dehydration series and CPD. For this we selected four Dipteran vector species: three mosquitoes *Aedes, Anopheles, Culex*, and one tsetse *Glossina* (specimen metadata in Supplementary Table S1). Specimens from each genus were selected across three decades (1930s, 50s and 70s) with the idea of assessing differences in DNA retrieval and morphological damage caused to samples of different ages. However, there was substantial variation within and between these species/decade sets, with samples from the 1930s and 50s occasionally containing longer DNA and higher yields than samples from the 70s (Supplementary Table S3, Supplementary Figure S10). This is likely due to initial specimen handling at collection and storage prior to being archived at the NHM, information which is very often missing from the pinned specimen labels.

A detailed evaluation of morphological changes post DNA extraction and CPD, as well as our internal score system detailing what level of damage we count as “pass” (key diagnostic features still preserved) or “fail”, is outlined in Supplementary Table S2. We noticed that the lowest level of morphological damage across all species was obtained with lysis buffer C (11 pass, 1 fail), with buffer A performing slightly worse (8 pass, 3 fail), and buffer G performing the worst (5 pass, 7 fail) (representative specimens across genera for each buffer showcased in Supplementary Figures S14-S17, with a closer look at an *Anopheles* lysis buffer C “pass” specimen in Figure 3A). When looking across the different genera regardless of lysis buffer, the most affected were *Anopheles* (3 pass, 5 fail), followed by *Aedes* (5 pass, 4 fail) and *Culex* (7 pass, 2 fail), while all nine *Glossina* specimens got a passing score.

**Fig. 3.**
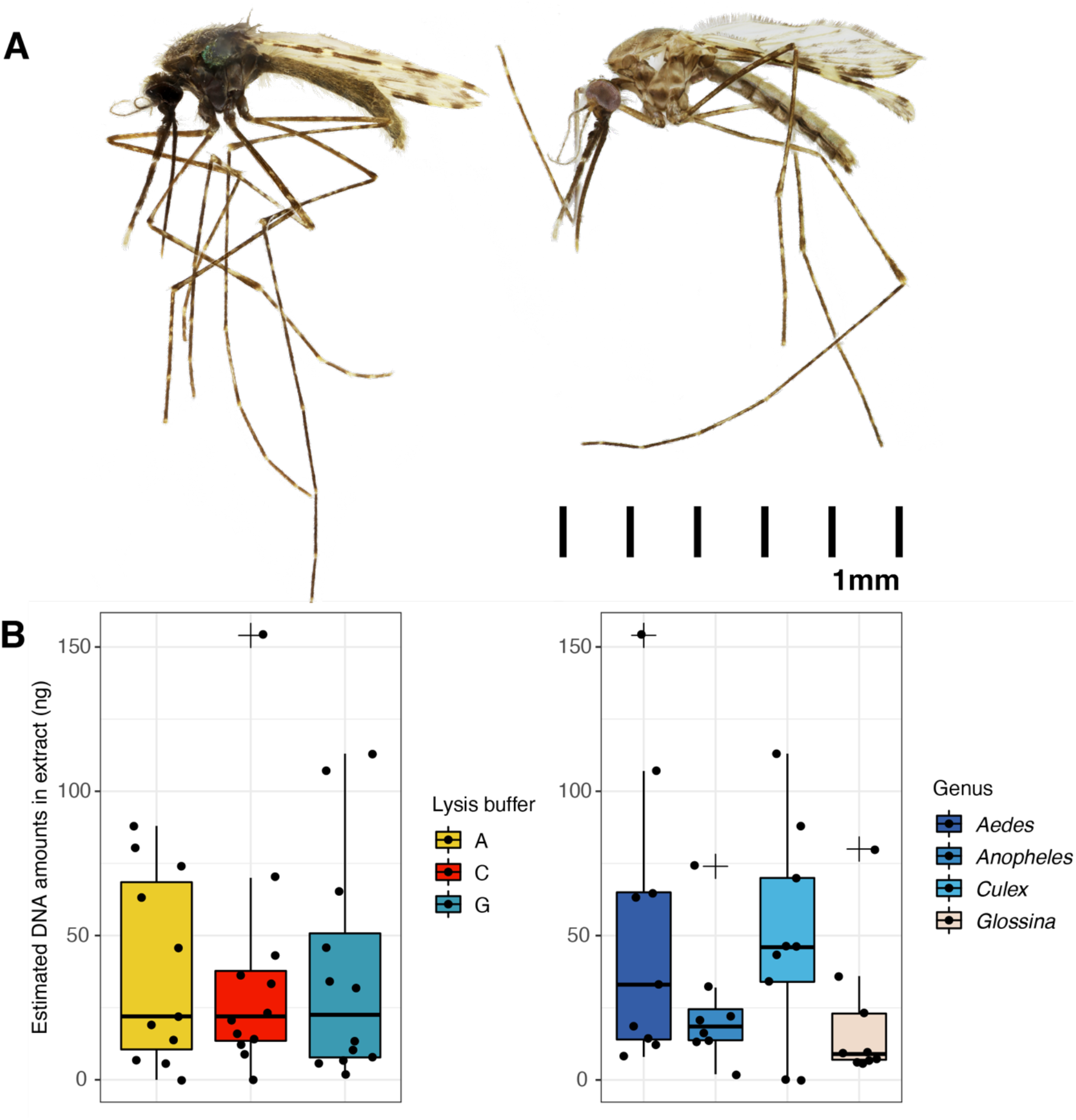
Lysis buffer efficiency across four Dipteran vectors. A) Focus stacked images of NHMUK010633504 (*Anopheles*) before (left) and after (right) a 2 h DNA extraction with lysis buffer C. This specimen sustained minimal morphological alterations after extraction. Adobe Photoshop was used for normalizing brightness and contrast, resizing (with millimetre scale attached), rotating and removing backgrounds. An overview of other representative specimens from all four genera and all three buffers is presented in Supplementary Table S2 and Supplementary Figures S14-S17. B) Estimated DNA yields in nanograms (ng) measured by Quant-iT™ PicoGreen™ dsDNA Assay Kit. Left is grouped by buffer, right is grouped by genus.

In terms of DNA retrieval, we noticed a similar release between the three buffers across all species, with slightly more DNA retrieved from *Aedes* and *Culex* compared to *Anopheles* and surprisingly *Glossina*, which had very low DNA yields for their size (Figure 3B). This might be due to extensive soft tissue degradation in the abdomen and thorax (specimens appeared to be hollow). It is also possible that larger samples such as *Glossina* require longer incubation times than 2 h in order for the buffer to penetrate deeper through the outer chitin layer. DNA length was also very variable between different species, decades, and lysis buffers (Supplementary Figure S10).

## Discussion

Historic museum collections are invaluable snapshots through space and time of populations that lived in a period prior to, during, and directly after extreme anthropogenic influences, such as the widespread use of insecticides in order to control populations of disease vectors (Forgash 1984; Kleinschmidt & Rowland 2019). With ever evolving methods for the retrieval and sequencing of old, fragmented and low yield DNA, primarily for ancient DNA research of fossil bones and teeth, we now can also apply and modify such methods to obtain genomic data from historic specimens (Staats et al. 2013; Gutaker et al. 2017; Parejo et al. 2020; Andrade Justi et al. 2021). In this paper we have focused on important human disease vectors, especially sub-Saharan malaria transmitting *Anopheles* mosquitoes. Over the course of three experimental setups we have evaluated the amount of DNA that can be retrieved from dried and pinned historic specimens seeking approaches that minimize the level of morphological damage caused during sample handling while maximizing the amount of informative genomic data we can obtain.

Across all of our experiments we have identified key points that are crucial in minimizing the level of morphological damage afflicted to the specimen during DNA extraction. These include rehydration of desiccated tissues, a very short lysis (2 h), and critical point drying soon after the lysis is performed to minimize damage from conventional air drying or prolonged exposure to ethanol. We also noticed that the initial mounting procedures, including the thickness of pins that pierce the specimen’s thorax and general pin placement, affects contact with liquids or plasticware that may degrade the specimen’s morphology. One of the biggest risks we identified was that if pins were placed too high up in the thorax, it led to decapitation or neck extension (Supplementary Figure S2). Fortunately, any larger body pieces such as limbs, the head or abdomen, can be collected and stored in capsules together with the specimen’s body since, although the specimen is no longer intact, no tissue is mechanically destroyed by grinding during the extraction procedure. We have also noticed that some surface structures on certain species tend to be more prone to damage during the extraction and drying procedure, such as bristles on the head and thorax, as those were lost across nearly all species presented in this paper (Supplementary Table S2). However, other key morphological features, such as scales across the body and limbs, including wings, were largely unaffected in all samples. Furthermore, even though 2 h seemed to work perfectly fine for most, shorter or longer incubation times might be required for some species, as seen for the very low DNA release for physically much larger *Glossina* samples.

In terms of DNA retrieval, we saw a lower total DNA yield compared to present-day samples extracted across all three used buffers, which was expected due to degradation of DNA through time (Supplementary Figure S11, present-day data from (Makunin et al. 2021)). Our samples were at the lower end of DNA yields compared to present-day samples (yields of 43±43 ng on average across all buffers and incubation times compared to 172±99 ng for present-day samples), as well as severely fragmented in most cases (Supplementary Figure S3, S6, S10). Because of that we modified our library preparation to accommodate for such short low yield inserts and noticed that less stringent library preparation approaches can be used compared to regular ancient DNA libraries. To make processing historic samples faster we opted for no shearing for samples that still contain slightly longer DNA, as well as SPRI bead purification post-ligation instead of column cleanups, the concentrations of which were adapted to fit a plate setup (2.2x post-ligation, two rounds of 1x post-indexing PCR). We have also shown that typical ancient DNA damage patterns are present in these samples, although as expected for DNA that is only decades old, at a much lower rate (up to 5% C>T at the 5’ end and G>A at the 3’ end, with 1-2% in the middle of the molecules, similar to what is observed for other historic tissues such as herbarium collections (Gutaker et al. 2017)) (Supplementary Figure S4, S7).

Our plex levels and sequencing depths (18-24 libraries on a single 75 PE HiSeq 2500 or 4000 lane, expected yields per lane 41 gigabases (Gb) and 43.75 Gb respectively, with insert sizes often below 150 bp) vary substantially from what is typically used for obtaining 30x coverage for present-day *Anopheles* specimens (36 libraries on three 150 PE HiSeq X10 lanes, expected yield for all three lanes is 330 Gb, with insert sizes often above 300 bp). While on average we retrieved about 2.6x mean nuclear coverage depth (from 0.4 to 9.3x), we were also able to retrieve 372x mean mitochondrial coverage (from 10 to 1,583x), and could assemble consensus mitochondrial genomes for all samples, including very low yield samples extracted with lysis buffers without any Proteinase K. Additionally, our historic samples have higher complexity than typical ancient DNA libraries, and the over-sequencing of PCR duplicates is fairly low (average duplication rate of 0.12±0.05 across sample libraries, Supplementary Table S3). This means there are still plenty of unique sequences present in each library, and additional sequencing can be performed to reach coverage levels of 20-30x which will facilitate genotyping and minimize potential biases caused by ancient DNA substitutions. As for the consensus mitochondrial genomes, we compared them to NCBI available genomes, some of which were published mitochondrial phylogenies of species in the *An. gambiae* complex (Beard et al. 1993; Peng et al. 2016; Hanemaaijer et al. 2018) and *An. funestus* complex (Hua et al. 2016; Peng et al. 2016; Jones et al. 2018; Liu et al. 2019; Small et al. 2020), and created maximum likelihood trees using MAFFT and FastTree incorporating both present-day samples and historic specimens (Supplementary Figure S8, S9). This helped in assessing that two samples were misidentified, and one could be mapped to the proper reference. The potential of creating mitochondrial genomes could help immensely with categorizing misidentified specimens in collections, especially species with less prominent vertical gene transfer and hybridization potential as malaria transmitting *Anopheles*. Even though low coverage, we also checked known insecticide resistance variants in present-day *An. gambiae* complex populations in the voltage-gated sodium channel gene (VGSC) (Clarkson et al. 2021), and found no evidence of the emergence of known insecticide resistance variants in our sequenced specimens (Supplementary Table S4). However, our dataset requires deeper sequencing and sequencing of more specimens to have the necessary resolution to assess the origin of insecticide resistance variants in more detail.

Across all our experiments we have noticed minimal difference between the three lysis buffers used, however lysis buffer C slightly outperforms the others when looking at the level of morphological damage after DNA extraction. As mosquitoes are quite fragile compared to other species of flies, more work is required in order to get a better understanding on what are the ideal extraction conditions (temperature, time) for different species to minimize morphological damage even further. We recommend doing an initial DNA extraction test on present-day samples (as already highlighted for *Anopheles* specimens in (Makunin et al. 2021)) to assess the level of morphological damage that might be caused by handling and exposure to lysis buffer, especially noting down changes in morphologically relevant traits such as protein-based pigments, which will likely be destroyed with proteinase based lysis buffers (Santos et al. 2018). A very important part of the process would also be the selection of samples from the start, as poorly preserved samples are likely to be further damaged in the whole process. Other optimizations could include streamlining the DNA purification process by using silica beads instead of columns, so the whole procedure could be performed in plates instead of single tubes (Rohland et al. 2018).

We hope our approach will further help with elucidating population structure changes in insect species, such as disease transmitting Diptera or those most affected by the climate crisis, and we will be able to study the changes observed in the genomes of present-day individuals in real time across the last century, while also preserving precious and limited historic pinned samples for future generations.

## Material and Methods

### Historic pinned specimen selection

Across all experiments we retrieved DNA from 87 pinned dry Diptera specimens from the NHM’s 2.5 million specimen Diptera collection (verbatim labels in Supplementary Table S1). Based on the labels accompanying each specimen these were classified as *Anopheles gambiae* (n = 20), *An. melas* (n = 10), *An. merus* (n = 2), *An. arabiensis* (n = 2), *An. funestus* (n = 26), *Aedes aegypti* (n = 9), *Culex pipiens* (n = 9) and *Glossina morsitans* (n = 9). The specimens were collected across a wide range of years and locations, from 1927 to 1988 and spanning 14 countries. Specimens were imaged both before and after DNA extraction using a Canon 5DSR with an Mp-E 65 mm stackshot rail for stacking. The wedge lights and specimen platform were custom built by the engineering department at the NHM. Eos Utility V.3, Helicon Remote, and Helicon Focus were used to create focus stacked images. Images were also taken at the Wellcome Genome Campus using a Hirox 3D digital microscope (before extraction photos of a few representative specimens in Supplementary Figure S12).

When working with historic specimens, similar to other ancient DNA work, it is critical to minimize the effects of modern day contaminants. Therefore, pinned historic Diptera samples were handled in laboratories where no present-day Diptera work was performed, especially avoiding post-PCR areas. During DNA extraction and purification buffers were prepared and handled inside UV decontaminated PCR cabinets, and most reagents (besides SDS and Proteinase K) and all DNA LoBind plasticware were decontaminated in a UV crosslinker 2x 45 min prior to use. Aliquots of purified DNA extracts were then transferred to post-PCR areas for quality control (concentration and fragment length measurements) as well as library preparation and sequencing. For each set we included several extraction blanks (tubes containing no sample DNA, just buffers) which were processed the same way and sequenced in the same pools at a lower fraction (Supplementary Table S3). We found no concerning sign of contamination with present-day DNA or cross-contamination with historic DNA in any of our blanks. Raw sequencing data for all specimen and blank libraries has been deposited in the European Nucleotide Archive (ENA) under study accession ERP129396, FASTQ IDs specified next to their corresponding NHM IDs in Supplementary Table S3.

### Initial assessment of DNA preservation in historic *Anopheles*

First we tested the efficiency of DNA retrieval from 10 pinned historic *An. gambiae* complex specimens using a minimally destructive low salt Proteinase K buffer described for insect tissue clarification prior to microscopy (consisting of 200 mM Tris pH 8.0, 25 mM EDTA pH 8.0, 250 mM NaCl, 0.5% SDS and 0.4 mg/ml Proteinase K as described in (Santos et al. 2018)), in this paper defined as “lysis buffer A”. Two samples at the most extreme ages (1938 and 1988) were removed from their label pins but left on their sample pins, placed into 2.0 ml DNA LoBind tubes and 200 *μ*l of buffer A was added to completely submerge the sample. However, due to plasticware static electricity, the samples were torn apart during handling. Therefore, for the remaining eight samples we performed tissue rehydration for 3 h at 37°C in a styrofoam box containing wet paper towels prior to DNA lysis. After rehydration samples were removed from their label pins and submerged into 200 *μ*l of buffer A. All samples regardless of rehydration were then incubated in the buffer at 37°C overnight. The next day, lysis buffer was transferred into new tubes, while samples were rinsed with 500 *μ*l 100% ethanol for 30 min and then air dried before returning to the NHM. We observed substantial tissue collapse caused by air drying (especially eyes, abdomens, and antennae), which led us to evaluate critical point drying (CPD) with liquid CO_2_ to restore volume. Air dried samples were rehydrated in 30% ethanol, and a serial ethanol dehydration was performed (20 min incubation in 30% - 50% - 70% - 90% - 3×100% ethanol) followed by CPD on a Baltec CPD 030, which successfully restored volume to collapsed tissues (Supplementary Figure S2).

The lysates were purified using a MinElute PCR Purification Kit silica column approach optimized for the purification of an EDTA-rich lysis buffer used in DNA extraction from ancient bones and teeth (Dabney et al. 2013) with a few modifications. We added 200 *μ*l of lysis buffer A to 2.0 ml DNA LoBind tubes containing UV treated 1.4 ml Qiagen binding buffer (PB) and 55 *μ*l 3 M sodium acetate (7x the buffer volume instead of the 5x volume recommended in the manufacturer’s protocol or the 10x volume recommended by (Dabney et al. 2013)). After the full volume of lysis buffer, PB and sodium acetate mixture was centrifuged through, columns were washed twice with 750 *μ*l Qiagen wash buffer (PE), dry spinned at maximum speed, and the elution of silica bound DNA was performed twice with 25 *μ*l of TET buffer (10 mM Tris pH 8.0, 1 mM EDTA pH 8.0, 0.05% Tween-20) for a total eluate volume of 50 *μ*l stored in 1.5 ml DNA LoBind tubes. Quality control of the final extracts was performed using a Quant-iT™ PicoGreen™ dsDNA Assay Kit for concentration estimates (Supplementary Table S3), and an Agilent Bioanalyzer High Sensitivity DNA Analysis chip for concentration and DNA fragment length measurements prior to library preparation (Supplementary Figure S3). In total, 14 *μ*l of the full extract volume was used for library preparation for the majority of samples, while for lower yield samples (NHMUK012805007, extraction blanks) 28 *μ*l was used instead.

### Adapting library preparation and sequencing for low yield short DNA inserts

For simplicity and easier streamlining we modified the NEBNext^®^ Ultra™ II DNA Library Prep Kit for Illumina^®^, a single tube double stranded sequencing library preparation kit already optimized and widely used at the Wellcome Genome Campus sequencing facilities (Bronner & Quail 2019), for low yield short insert DNA extracts. Based on the initial quality control evaluation, six samples (1930s-60s) had lower yields and short DNA, while four samples from the 1980s had higher yields and longer DNA. For the 1980s extracts at the start of library preparation the volume was split in half, with one half undergoing Covaris shearing prior to library preparation, and the other going into library preparation without shearing. For all other extracts the post-adapter ligation reaction was split in half, one half was purified using silica columns (MinElute PCR Purification Kit) and the other using 3x SPRI beads (Beckman Coulter™ Agencourt AMPure XP). Indexing PCR, which adds 8 bp sample-specific indices on both ends of library molecules, was performed using KAPA HiFi HotStart ReadyMix following the manufacturer’s protocol for a total of 10 PCR cycles. Unfortunately, this was performed using a version of Kapa HiFi that cannot recognise uracils, and therefore the observed ancient DNA specific substitution rates do not reflect the actual deamination rates in these samples (Supplementary Table S3, Supplementary Figure S4). After indexing PCR, amplified libraries were purified with a combination of 3x SPRI, silica columns and 1x SPRI beads until no detectable adapter dimer peak was visible and validated on an Agilent Bioanalyzer High Sensitivity DNA Analysis chip. All sample libraries were then pooled equimolarly, with extraction blanks pooled at a 1:10 molar ratio compared to sample libraries, and sequenced on one lane of an Illumina HiSeq 2500 System 75 PE with two additional 8 bp index reads.

After sequencing, reads were split into cram files for each library based on matching 8 bp tags, converted into FASTQ files and processed using the ancient DNA analysis pipeline EAGER (Fellows Yates et al. 2021) (nextflow version 20.10.0, EAGER last modified December 2020). The following parameters were used: adapter sequence trimming (forward AGATCGGAAGAGCACACGTCTGAACTCCAGTCACNNNNNNNNATCTCGTATGCCGTCTTCTGCTTG, reverse AGATCGGAAGAGCGTCGTGTAGGGAAAGAGTGTNNNNNNNNGTGTAGATCTCGGTGGTCGCCGTATCATT), aligning to the *Anopheles gambiae* reference genome (AgamP4) using bwa mem, merging overlapping reads (with default minimum 11 bp overlap), not filtering unmerged reads (longer inserts in younger samples), performing DamageProfiler for a summary of ancient DNA characteristics (5’ C>T and 3’ G>A substitutions, read length in base pairs), removing PCR duplicates and unaligned reads for final bam files. Additionally, samtools coverage was used on the final filtered bam files to get details on nuclear and mitochondrial depth of coverage, as well as the number of covered bases. Final bam files for libraries prepared using different strategies (purification with beads vs columns, unsheared vs sheared) for each of the 10 samples were merged, fragments mapping to the mitochondrial genomes were extracted and a consensus mitochondrial genome was created from each sample using bcftools mpileup. Sequencing summary statistics for each library are presented in Supplementary Table S3.

### Testing different lysis conditions on morphological damage versus DNA retrieval

We next tested different lysis buffer and incubation times on the efficiency of DNA release and the level of morphological damage on different *Anopheles* species. Samples morphologically designated as *Anopheles gambiae* (n = 8), *An. melas* (n = 8), and *An. funestus* (n = 26) were extracted using three lysis buffers, whose performance was assessed in detail on present-day *Anopheles* species in (Makunin et al. 2021): lysis buffer A (Santos et al. 2018), lysis buffer C (simplified A: 200 mM Tris pH 8.0, 25 mM EDTA pH 8.0, 0.05% Tween-20 and 0.4 mg/ml Proteinase K), and lysis G (simplified (Gutaker et al. 2017): 10 mM Tris pH 8.0, 10 mM EDTA pH 8.0, 5 mM NaCl, 0.05% Tween-20 and 0.4 mg/ml Proteinase K). Samples were rehydrated 2-3 h at 37°C, removed from their respective label pins and submerged in 200 *μ*l of buffer A, C or G overnight (34 samples) or 2 h (8 samples) in an oven at 37°C. Four samples (2 *An. gambiae* and 2 *An. melas*) were incubated in lysis buffers C and G without the addition of Proteinase K to assess the level of DNA retrieval and morphological damage with minimal tissue clarification. After incubation, lysis buffer was transferred to new tubes, and the samples were ethanol dilution washed (500 *μ*l 30 - 50 - 70% ethanol, 20 min each) and either stored in 70% ethanol or fully washed with an additional 90% and 3×100% (stored in final 100% ethanol volume), before returning to the NHM for CPD. Unfortunately, due to a CPD instrument malfunction there was still substantial morphological damage as a result of the lysis process and/or prolonged desiccation in high ethanol concentrations, which causes tissue collapse.

Lysates were purified using the same MinElute silica column approach, as we confirmed in a ladder experiment using both a short (Thermo Scientific GeneRuler Ultra Low Range DNA Ladder) and long (Thermo Scientific GeneRuler 1 kb Plus Ladder) DNA ladder that we are able to retrieve DNA fragments in the range of 25-10,000 bp using this modified MinElute approach (Supplementary Figure S5). DNA extracts were again evaluated using a Quant-iT™ PicoGreen™ dsDNA Assay Kit and Agilent Bioanalyzer High Sensitivity DNA Analysis chip or Agilent TapeStation High Sensitivity D5000 ScreenTape System (Supplementary Table S3, Supplementary Figure S6), and a total of 24 *μ*l was used to prepare libraries. Illumina libraries were prepared using the same NEBNext^®^ Ultra™ II DNA Library Prep Kit following the simplified approach identified in our initial experiment (no prior shearing, purification post-ligation with either 2.5x or 2.2x SPRI beads). For indexing PCR we used the KAPA HiFi HotStart Uracil+ ReadyMix PCR Kit in order to get correct deamination patterns. We used Bioanalyzer or TapeStation concentration values to estimate the optimal number of PCR cycles for each library: *An. gambiae* complex samples <10 ng - 12 cycles, 10-40 ng - 10 cycles, >40 ng - 7 cycles; *An. funestus* all samples 9 cycles. Post-PCR libraries were purified using one round of 1.2x SPRI beads (*An. gambiae* complex samples) or two rounds of 1x SPRI beads (*An. funestus* samples) to remove the majority of adapter dimers, libraries were checked on an Agilent TapeStation D5000 ScreenTape System, equimolarly pooled into three different pools (*An. gambiae* and *An. melas, An. funestus* overnight, *An. funestus* 2 h), again with a smaller proportion for extraction blanks in each, and the pools were sequenced on three lanes of an Illumina HiSeq 4000 System 75 PE with two additional 8 bp index reads. Summary statistics were prepared using EAGER with the same settings as previously described, with *An. gambiae* and *An. melas* samples being mapped to AgamP4 and *An. funestus* samples being mapped to the *An. funestus* nuclear (AfunF3) and mitochondrial (NC_038158.1) references. After our initial processing and mitochondrial DNA assembly, one of the *An. gambiae* samples (NHMUK010633485) showed a mitochondrial genome full of N-stretches which grouped with other *An. funestus* samples, and this sample was then mapped to the *An. funestus* reference instead.

### Morphological damage assessment across different Diptera species

In our final experiment we examined a wider range of Diptera disease vector species to evaluate the level of morphological damage caused by handling, DNA lysis and the drying procedure, as well as the amount of DNA that can be retrieved with a 2 h incubation in the previously tested lysis buffers. For this we selected 35 samples in total, morphologically designated as *Aedes aegypti* (n = 9), *Anopheles gambiae* (n = 8), *Culex pipiens* (n = 9), and *Glossina morsitans* (n = 9), with three samples each from three different decades (1930s, 50s and late 60s/early 70s defined as 70s henceforth). Samples were rehydrated for 3 h at 37°C, split into three (one for each decade) and incubated in 200 *μ*l of lysis buffer A, C or G for 2 h, except for *Glossina* which due to their size had to be incubated in 1 ml of lysis buffer in a tissue culture plate instead of individual 2.0 ml DNA LoBind tubes. After lysis all samples were rinsed with 500 *μ*l 30% ethanol, stored in 500 *μ*l 50% ethanol, and shipped to the NHM for CPD and imaging. Again, for *Glossina* samples volume was increased to 1 ml for the first wash and 2 ml storage in a 5.0 ml SafeLock tube due to their size. Lysis buffer was purified using the same modified MinElute silica column method. For *Glossina* we initially purified only 200 *μ*l, however not enough DNA was detected in this fraction and the remaining 800 *μ*l were purified and used for quality control assessment instead (increased volumes of PB to 5.6 ml and sodium acetate to 220 *μ*l). DNA extracts from all four species were evaluated using a Quant-iT™ PicoGreen™ dsDNA Assay Kit and Bioanalyzer High Sensitivity DNA Analysis chip or Agilent TapeStation High Sensitivity D5000 ScreenTape System (Supplementary Table S3, Supplementary Figure S10). As the goal of this experiment was primarily to compare morphological damage, we do not present library preparation and sequencing for this experiment.

## Acknowledgments

This research was funded by the Wellcome Trust Grant [206194] (which also supports M.K.N.L. and A.M); the London Natural History Museum; and the European Molecular Biology Laboratory. P.K. has been supported by the EMBL-EBI/Wellcome Sanger Institute Post-Doctoral Fellowship Programme (ESPOD). For the purpose of Open Access, the authors have applied a CC BY public copyright licence to any Author Accepted Manuscript version arising from this submission. The authors would like to thank the staff of the Wellcome Sanger Institute Scientific Operations for their contribution to library preparation and sequencing. We are also very grateful to members of the Lawniczak and Flicek Research groups for constructive discussions and comments during each step of the project, as well as Alex Ball and Innes Clatworthy (Imaging and Analysis Centre at the NHM core Research Laboratories) for help with the critical point drying process. And lastly, we are extremely thankful to the entomologists who decades ago collected and determined the Diptera specimens used in this study: Burtt, Buxton, Cattlin, Coetzee, Garnham, Gibson, Gillies, Harbach, Hunt, Knight, Leeson, Lewis, Mackie, Slater, Stymes, Surtees, Sutton, Vincent, and all supporting fieldwork and collection assistants whose names did not fit on specimen labels.

## Author Contributions

P.K., E.M., P.F. and M.K.N.L designed the project. P.K., E.M. and M.M. performed the experiments. P.K., E.M. and A.M. analysed the data. P.K., E.M. and M.K.N.L. wrote the manuscript with input from M.M, A.M. and P.F.

## Conflict of Interest

The authors declare no potential conflicts of interest.

## Data Availability Statement

Raw sequencing data for all specimen and blank libraries is available under ENA study accession ERP129396.

## Notes

https://www.ebi.ac.uk/ena/browser/view/PRJEB45310

### Competing Interest Statement

The authors have declared no competing interest.

