## Supplementary Information for "A minimally morphologically destructive approach for DNA retrieval and whole genome shotgun sequencing of pinned historic Dipteran vector species"

**This file includes:**

Supplementary Figures S1-S17
Supplementary Table S4

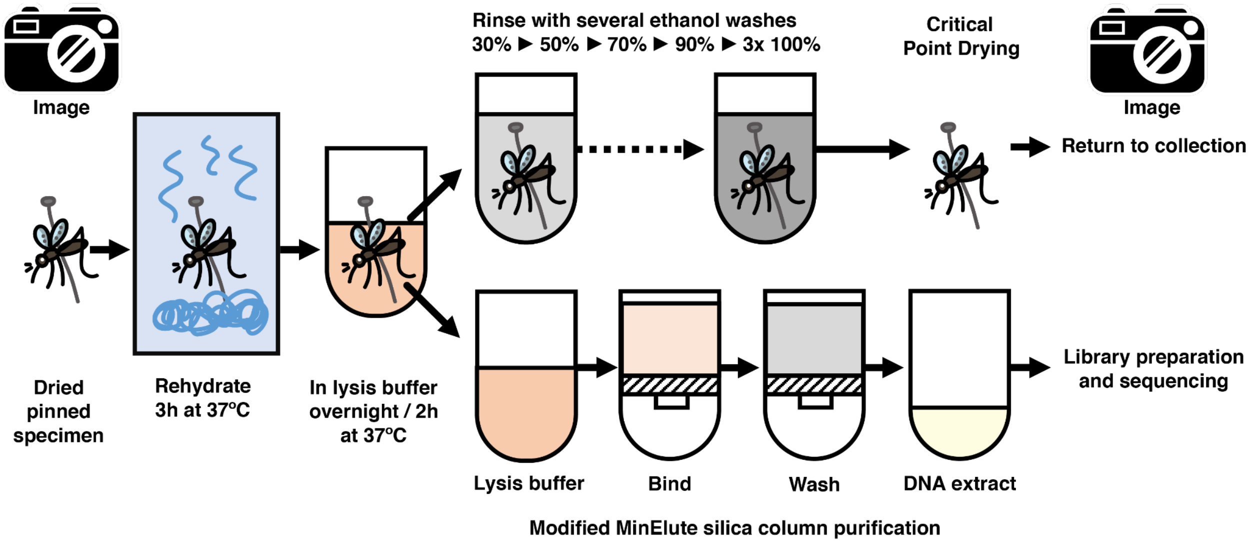


**Supplementary Figure S1.** Schematic view of our minimally destructive DNA extraction approach for historic pinned Diptera specimens. We recommend taking high resolution images of the specimens both at the first and last step.

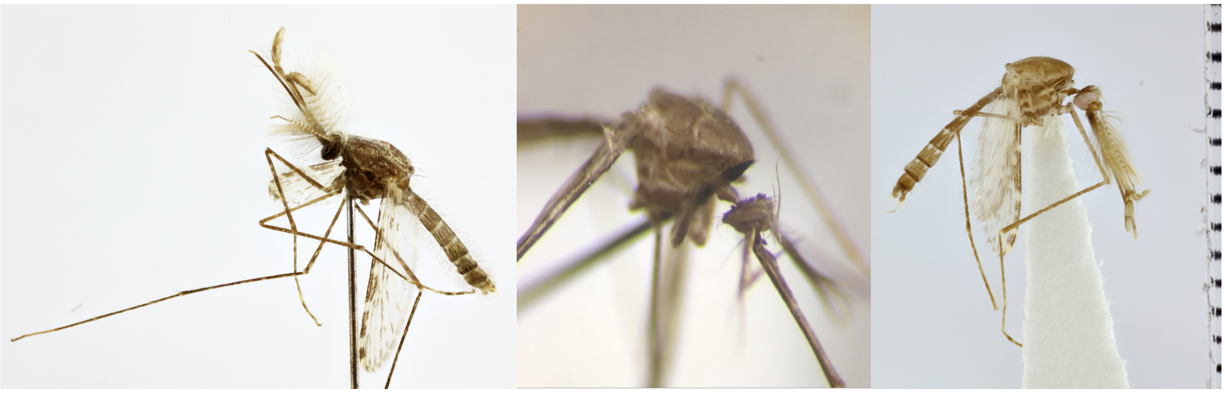

**Supplementary Figure S2.** Initial Critical Point Drying (CPD) assessment using NHMUK012805008 (*Anopheles melas* male specimen). **Left panel**: high resolution image of specimen pre-extraction. **Middle panel**: specimen after DNA extraction and air drying showcasing eye pigment loss, neck extension and 3D structure tissue collapse. **Right panel**: sample with restored 3D structure after CPD.

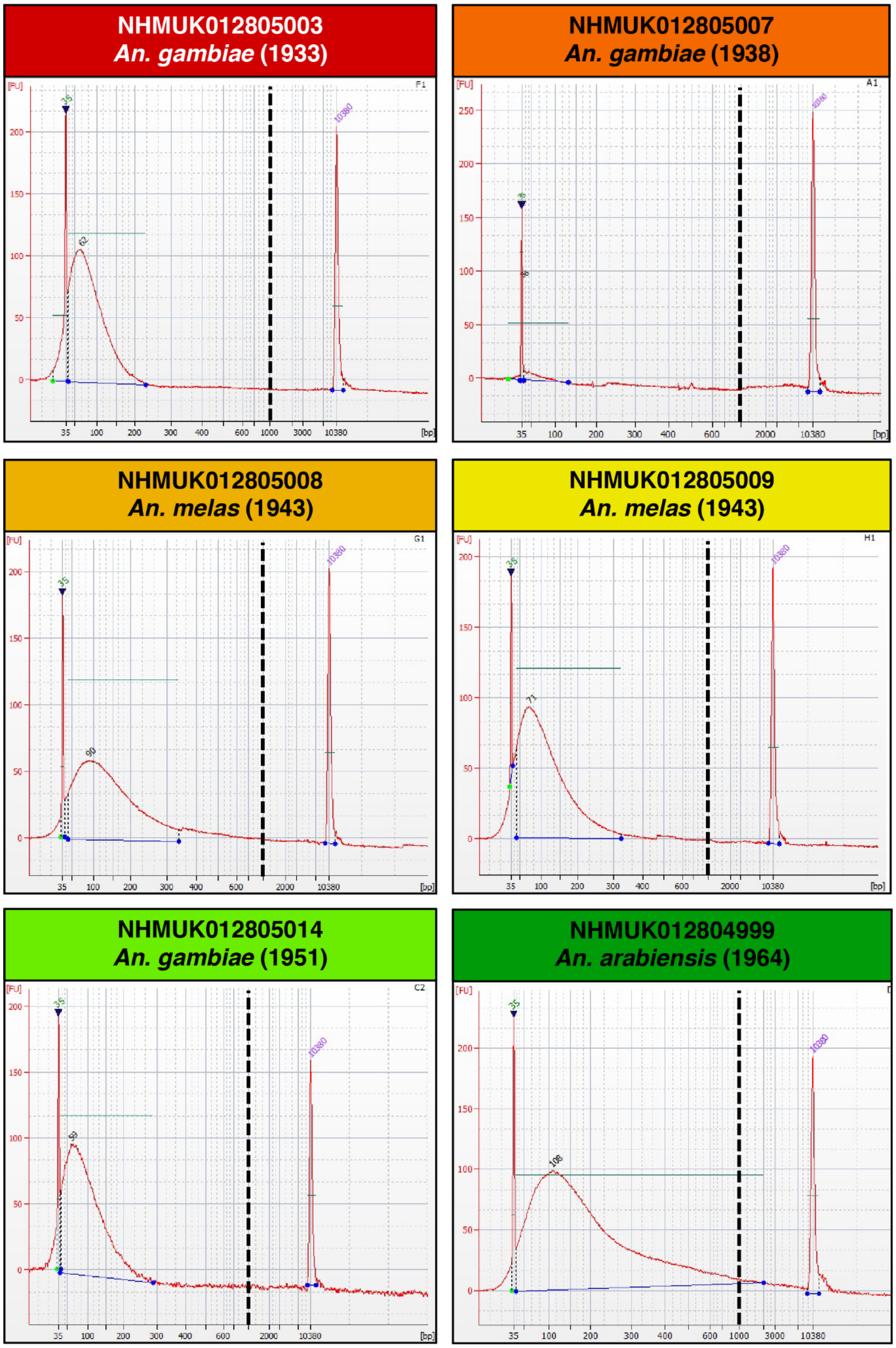


**Supplementary Figure S3.** Agilent Bioanalyzer High Sensitivity DNA Analysis Chip runs of DNA extracts prepared using lysis buffer A and MinElute silica column purification from 10 historic *An. gambiae* complex specimens (1930s-80s). A dashed vertical line is placed at the 1,000 bp mark on each panel as scales differ.

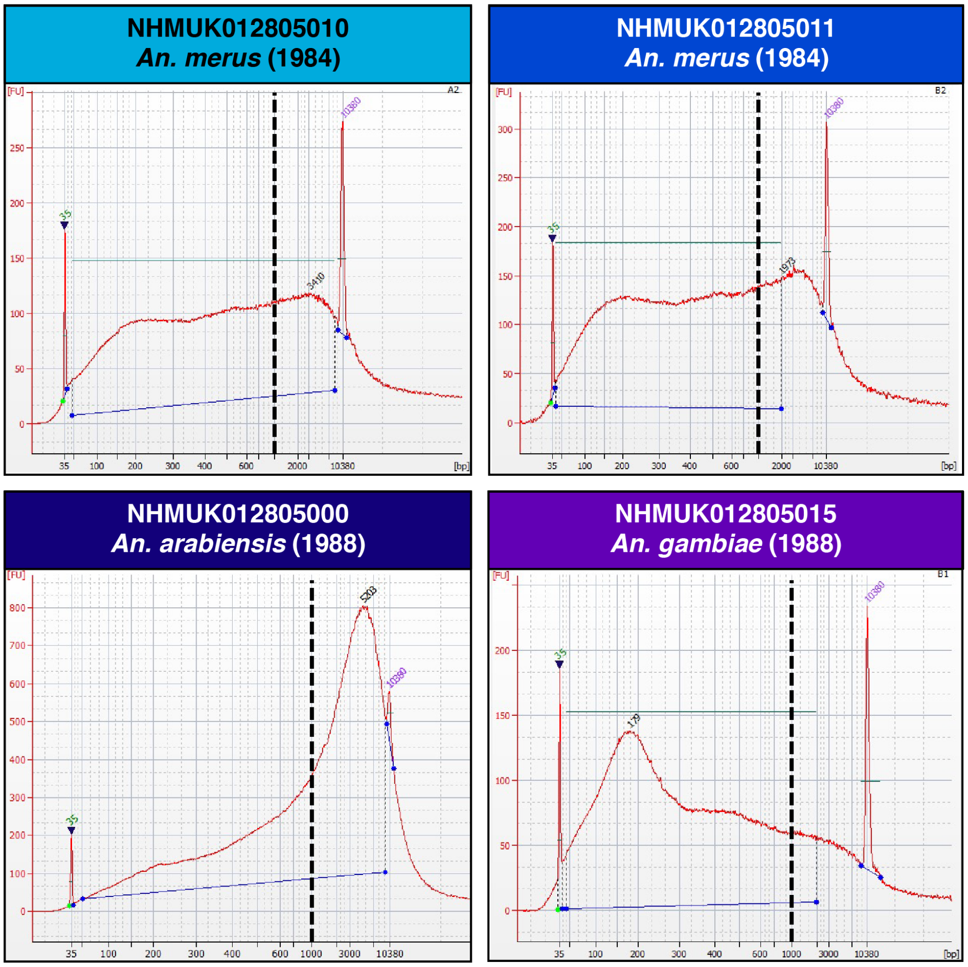


**Supplementary Figure S3. - continued**

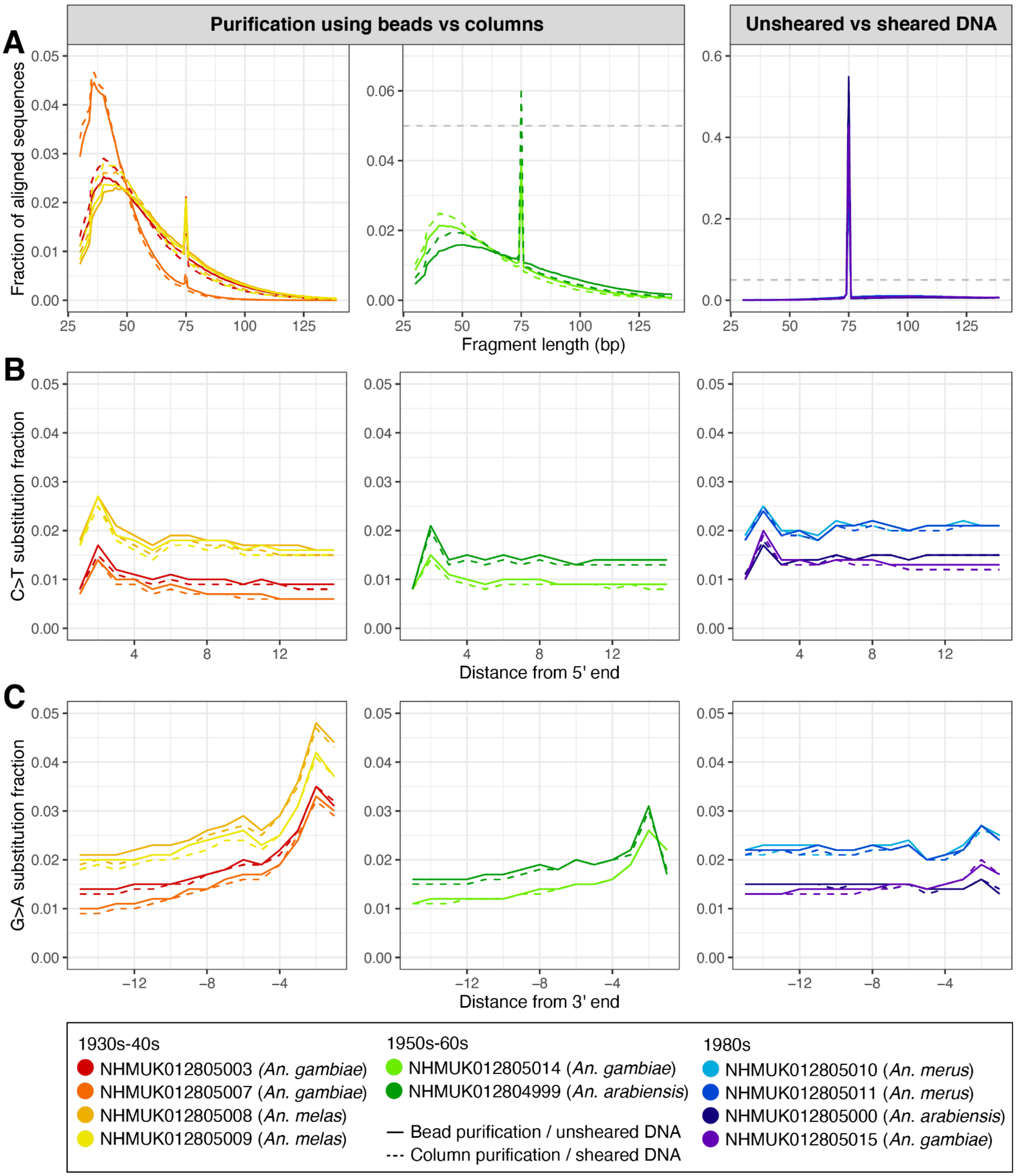


**Supplementary Figure S4.** Ancient DNA characteristics of 10 historic *An. gambiae* complex specimens. Dashed lines represent alternative library preparation strategies (left and middle panels: silica column purification post library preparation, right panels: DNA extracts sheared prior to library preparation). **A)** Size distribution of aligned DNA fragments. Samples with a higher fraction of 75 bp reads contain more DNA inserts >139 bp. The horizontal dashed line denotes the y axis limit from the first panel (0.05). **B)** C>T substitution fraction at 5' end. **C)** G>A substitution fraction at 3' end. As the polymerase used in indexing PCR does not recognise uracils, the 5' C>T substitution patterns are lower than expected, while the 3' G>A are more reliably represented since this substitution occurs during library preparation pre-PCR.

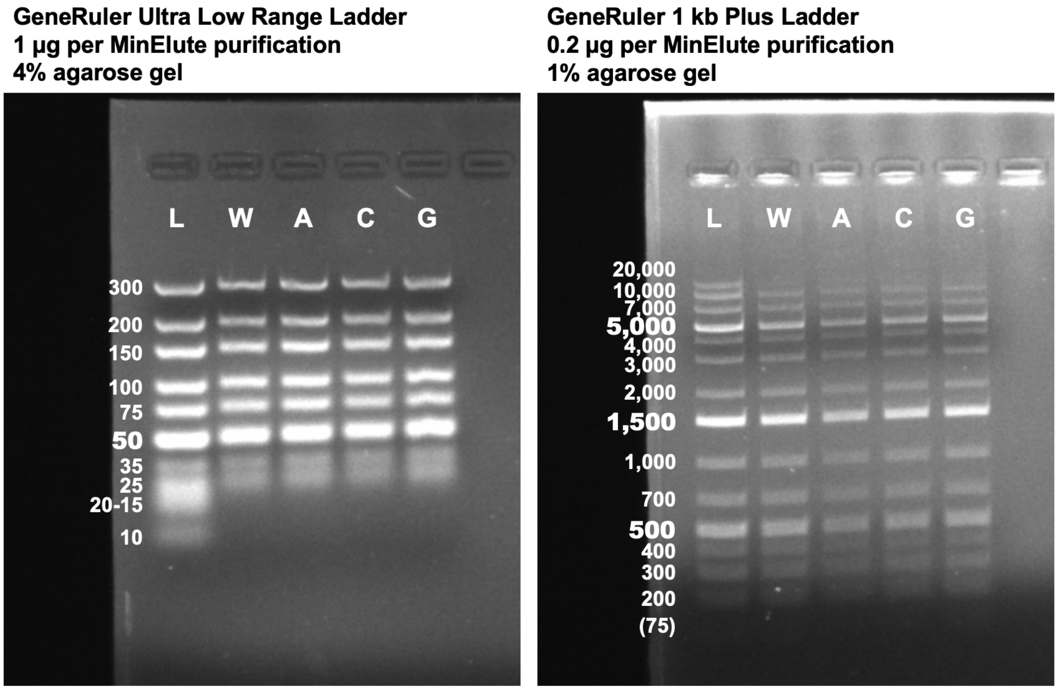


**Supplementary Figure S5.** Efficiency of DNA retrieval from various buffers using a modified MinElute silica column based DNA extraction. In short, 2 µl of ladder was added to 50 µl of water or lysis buffer and purified using the MinElute procedure described in Materials and Methods. **Left panel**: Thermo Scientific GeneRuler Ultra Low Range DNA Ladder ran on 4% agarose gel. **Right panel**: Thermo Scientific GeneRuler 1 kb Plus Ladder ran on 1% agarose gel. (L) ladder without purification, ladder purified from (W) water, (A) lysis buffer A, (C) lysis buffer C, (G) lysis buffer C. Agarose gels photographed using an Omega Fluor™ Gel Documentation System machine and labelled using Microsoft PowerPoint with minimal post-processing (gel crop).

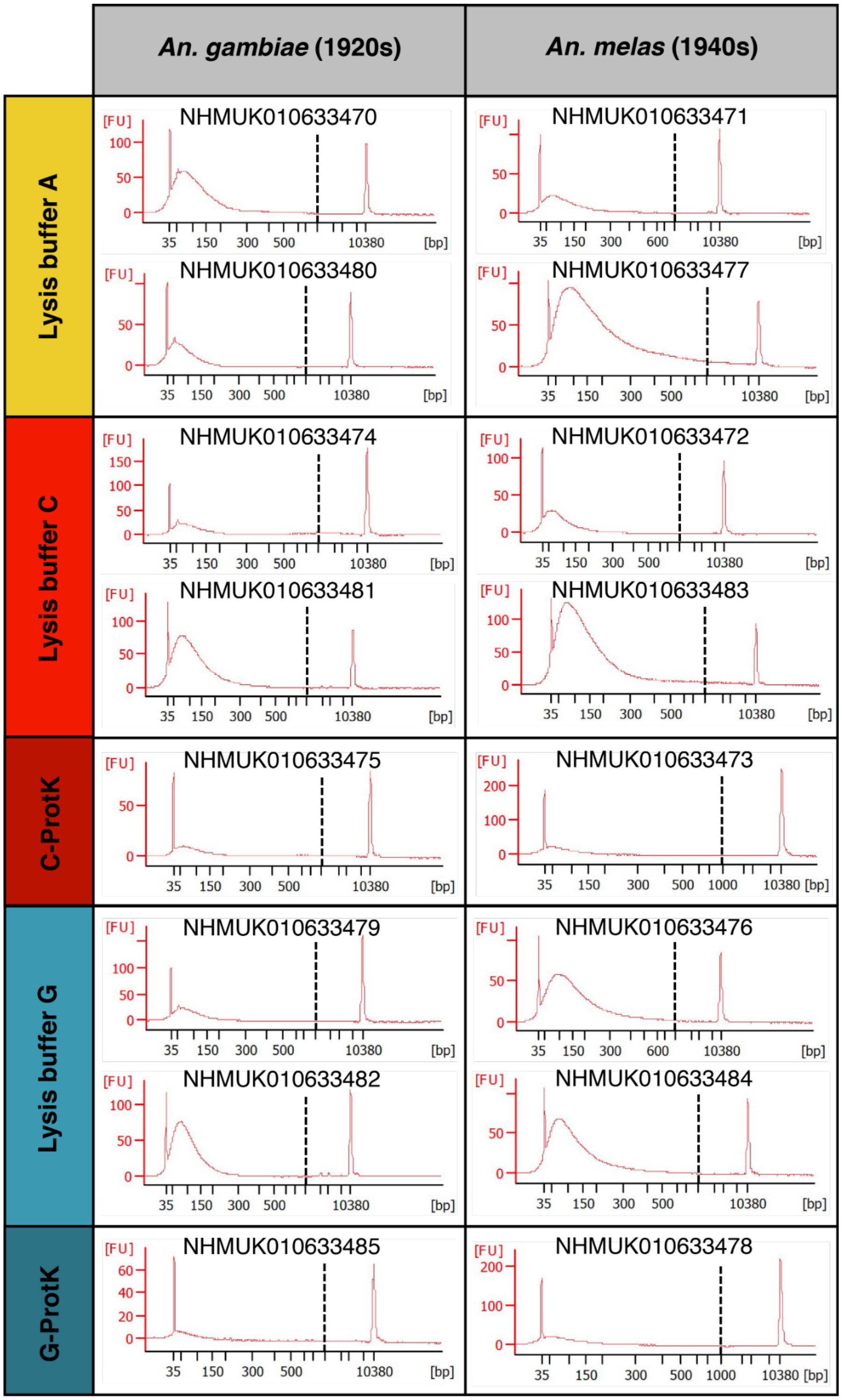


**Supplementary Figure S6.** Agilent Bioanalyzer High Sensitivity DNA Analysis Chip and Agilent TapeStation High Sensitivity D5000 ScreenTape runs of DNA extracts prepared using different lysis buffers and MinElute silica column purification from 8 *An. gambiae* from the 1920s, 8 *An. melas* from the 1940s, and 26 *An. funestus* from the 1930s (overnight and 2 h lysis incubation). The different lysis buffers used are noted in color on the y axis. A dashed vertical line is placed at the 1,000 bp mark on each panel as scales differ.

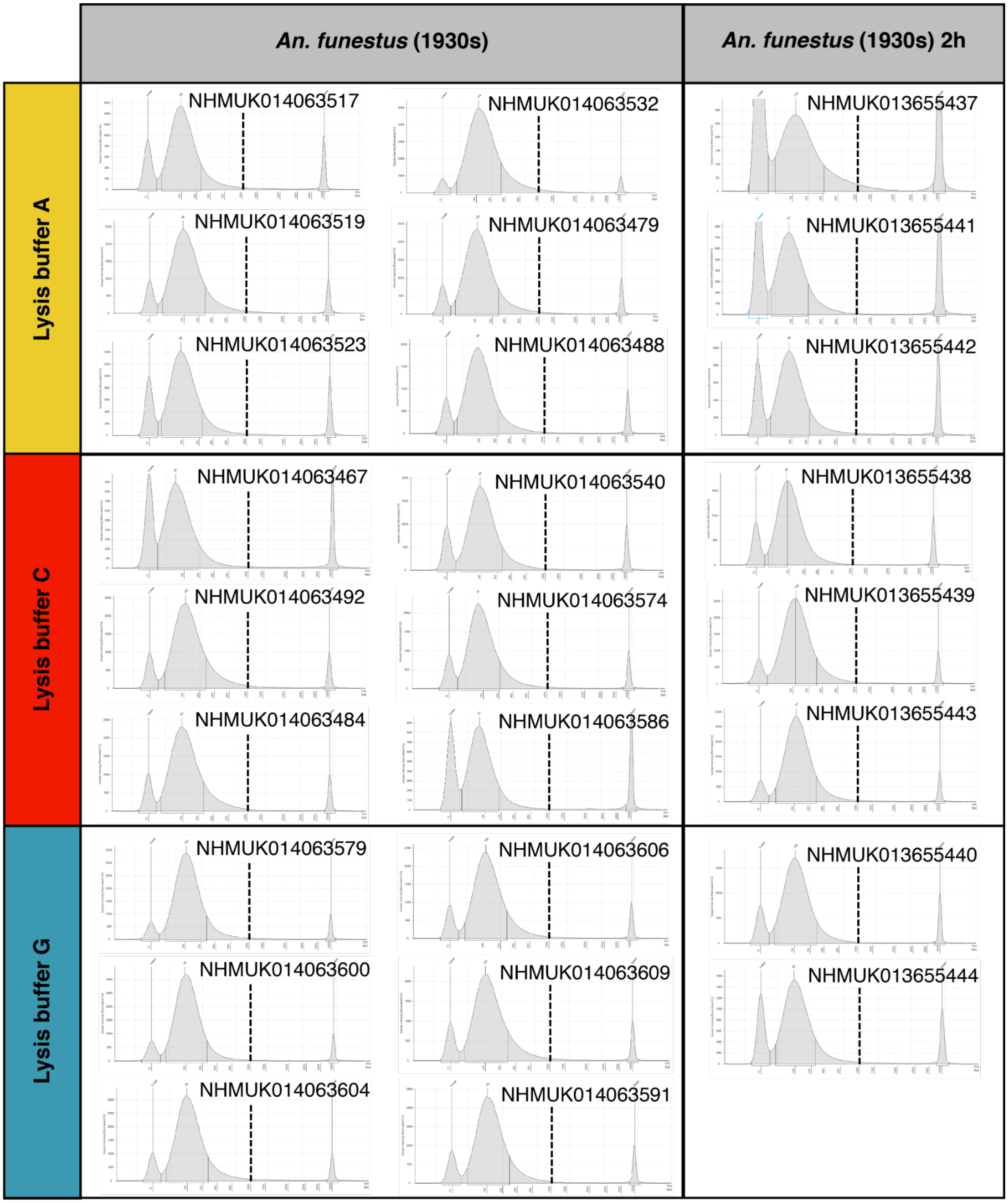


**Supplementary Figure S6. - continued**

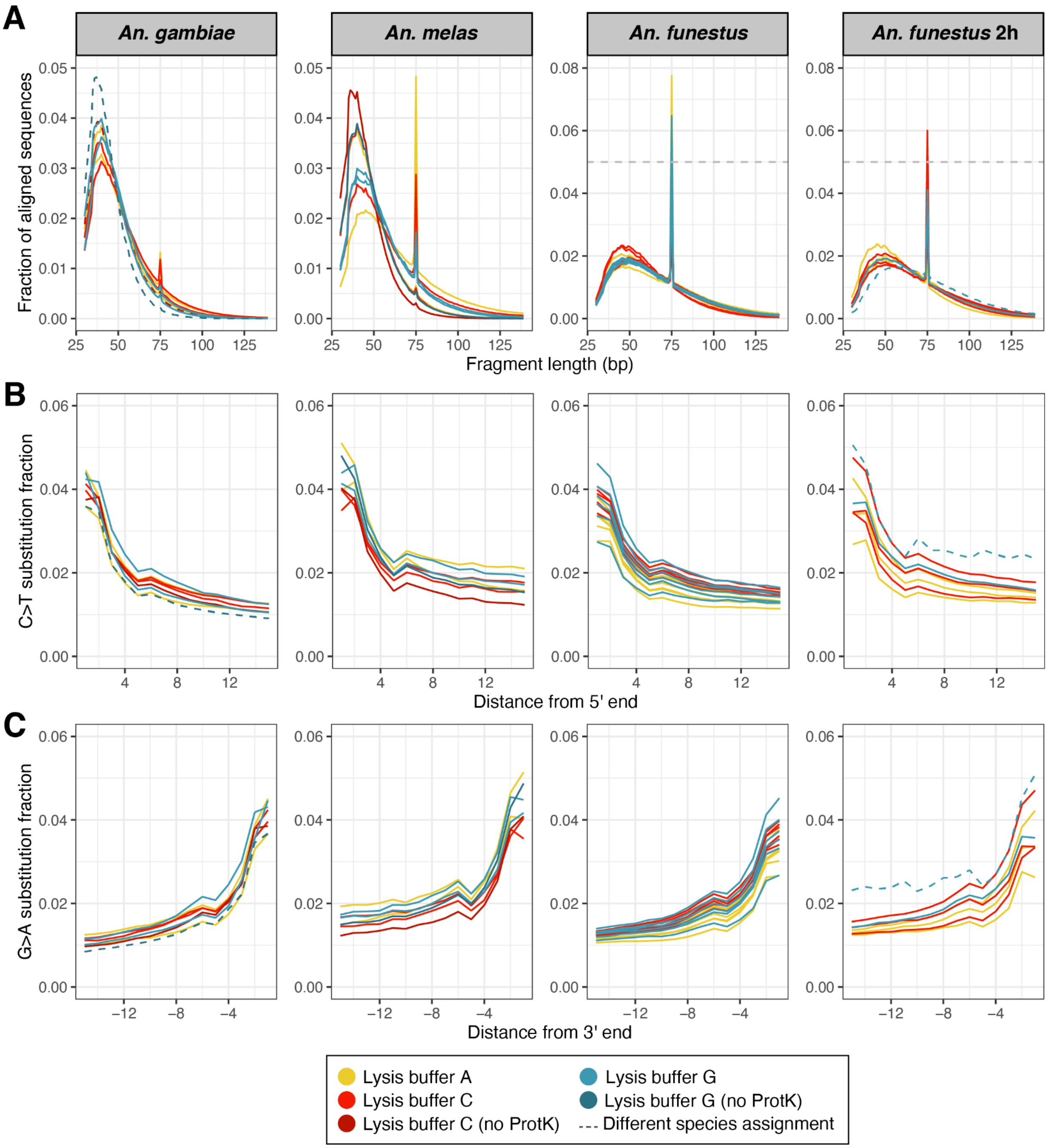


**Supplementary Figure S7.** Ancient DNA characteristics of 8 *An. gambiae* (1920s), 8 *An. melas* (1940s) and 26 *An. funestus* (1930s, extracted overnight or 2 h). Dashed lines represent samples that after sequencing were found to be different species from the ones they were stored as at the NHM collection (one *An. gambiae* is *An. funestus*, one *An. funestus* is *An. rivulorum*). **A)** Size distribution of aligned DNA fragments. Samples with a higher fraction of 75 bp reads contain more DNA inserts >139 bp. During library preparation the *An. funestus* samples were purified with less stringent SPRI bead ratios (Supplementary Table S3), resulting in an increased retrieval of longer sequences. The horizontal dashed line denotes the y axis limit from the first two panels (0.05). **B)** C>T substitution fraction at 5' end. **C)** G>A substitution fraction at 3' end. In these sets we used a polymerase that can recognise uracils, therefore both the 5' C>T and 3' G>A substitution patterns are as expected for aged DNA.

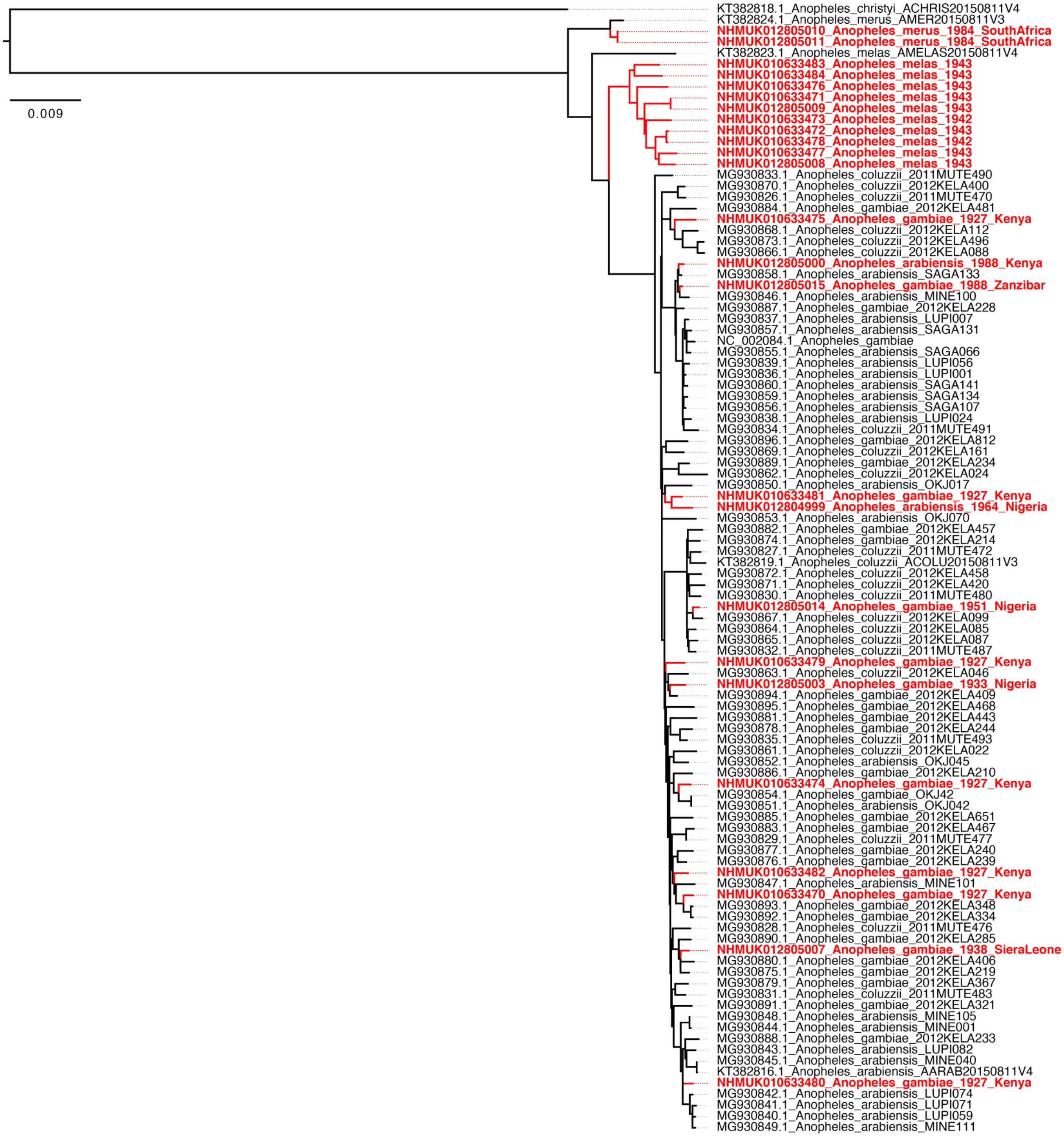


**Supplementary Figure S8.** Maximum likelihood mitochondrial genome phylogeny of samples falling into the *An. gambiae* complex. Historic specimens in red, NCBI available genomes (Beard et al. 1993; Peng et al. 2016; Hanemaaijer et al. 2018) in black.

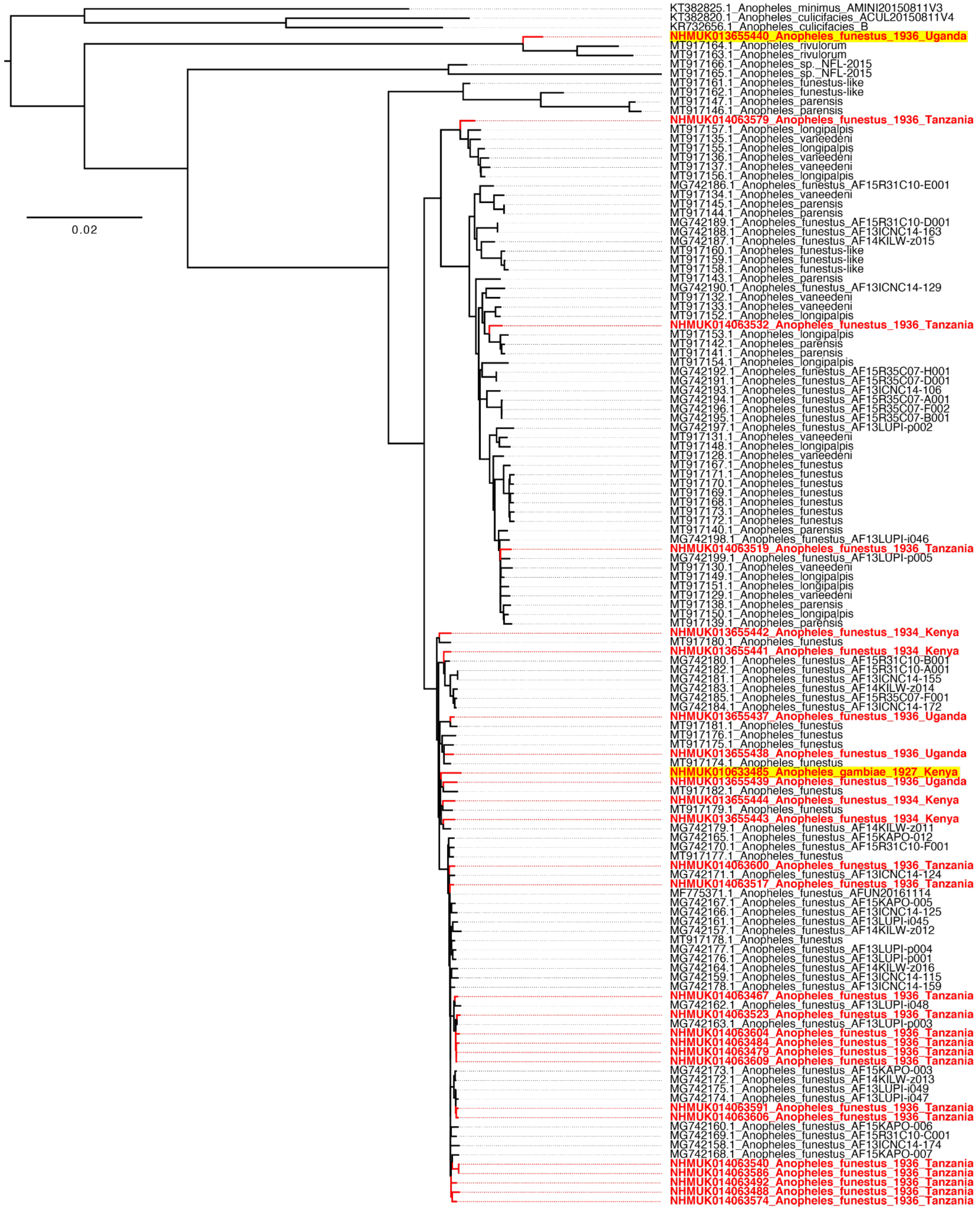


**Supplementary Figure S9.** Maximum likelihood mitochondrial genome phylogeny of samples falling into the *An. funestus* complex. Historic specimens in red, NCBI available genomes (Hua et al. 2016; Peng et al. 2016; Jones et al. 2018; Liu et al. 2019; Small et al. 2020) in black. Samples that do not match their original morphological assessment are highlighted in yellow.

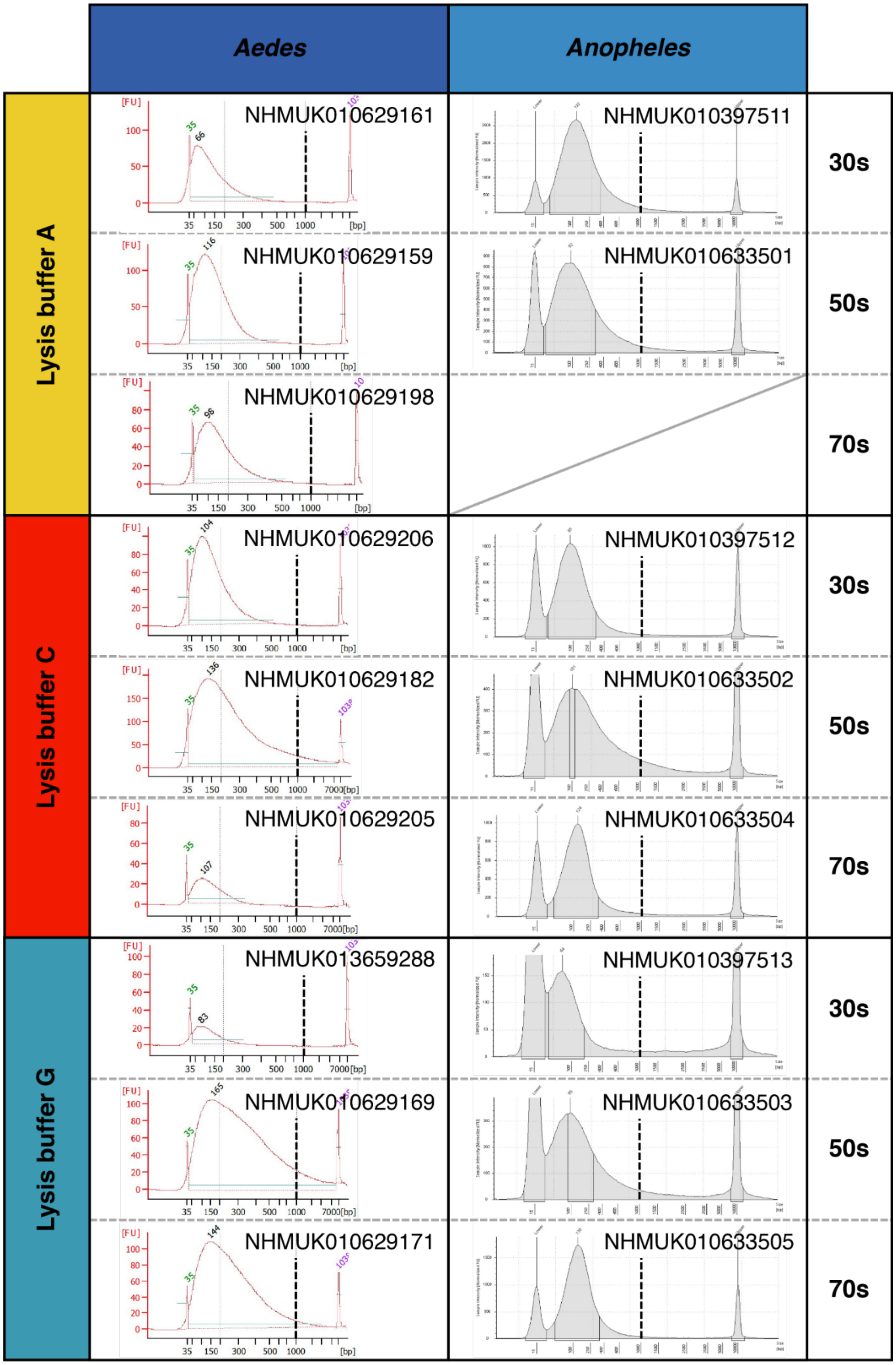


**Supplementary Figure S10.** Agilent Bioanalyzer High Sensitivity Chip and Agilent TapeStation High Sensitivity D5000 ScreenTape runs of DNA extracts prepared using different lysis buffers and MinElute silica column purification from four Diptera genera (columns) split by lysis buffers (left) and collection decade (right). A dashed vertical line is placed at the 1,000 bp mark on each panel as scales differ.

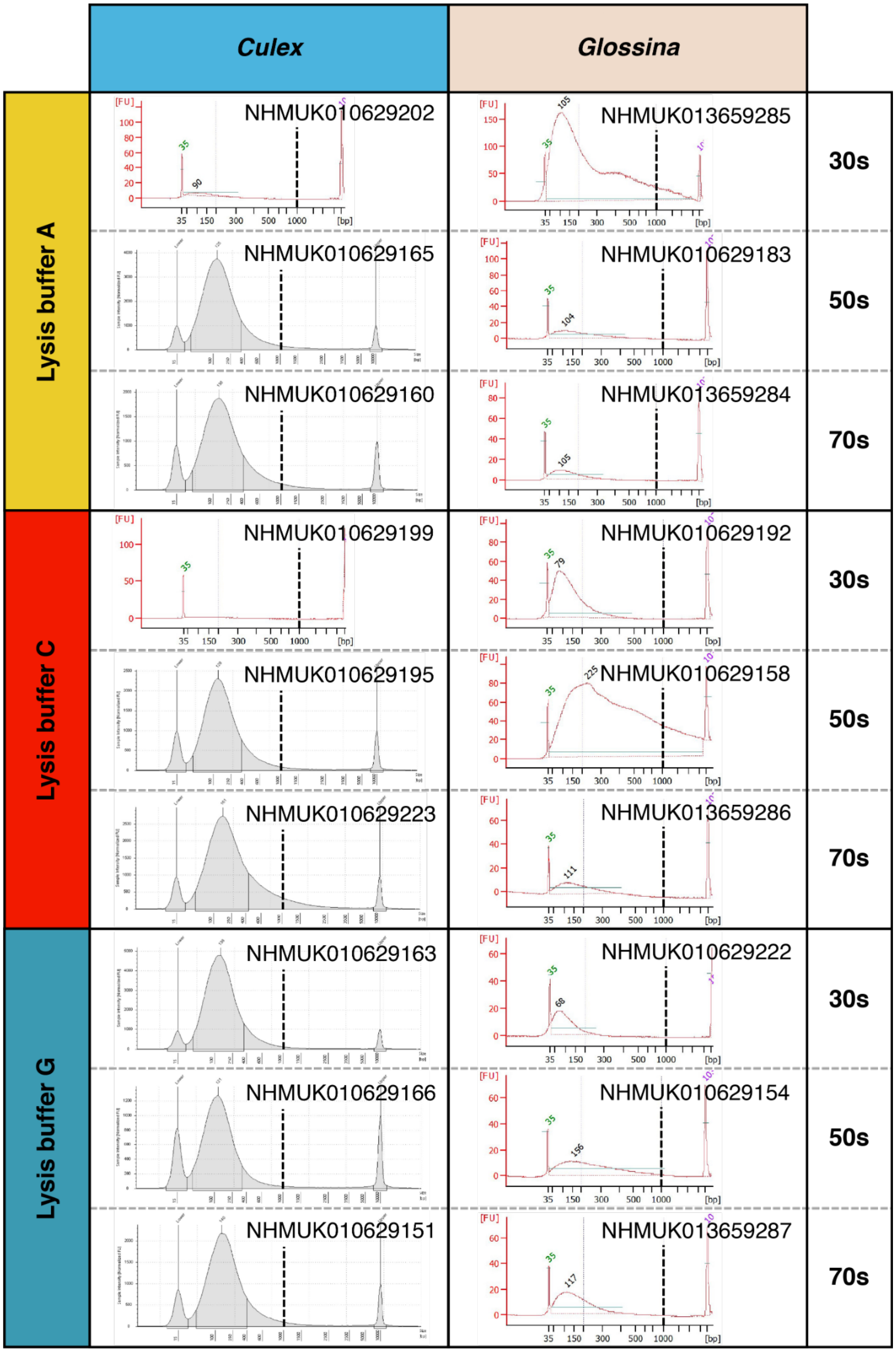


**Supplementary Figure S10. - continued**

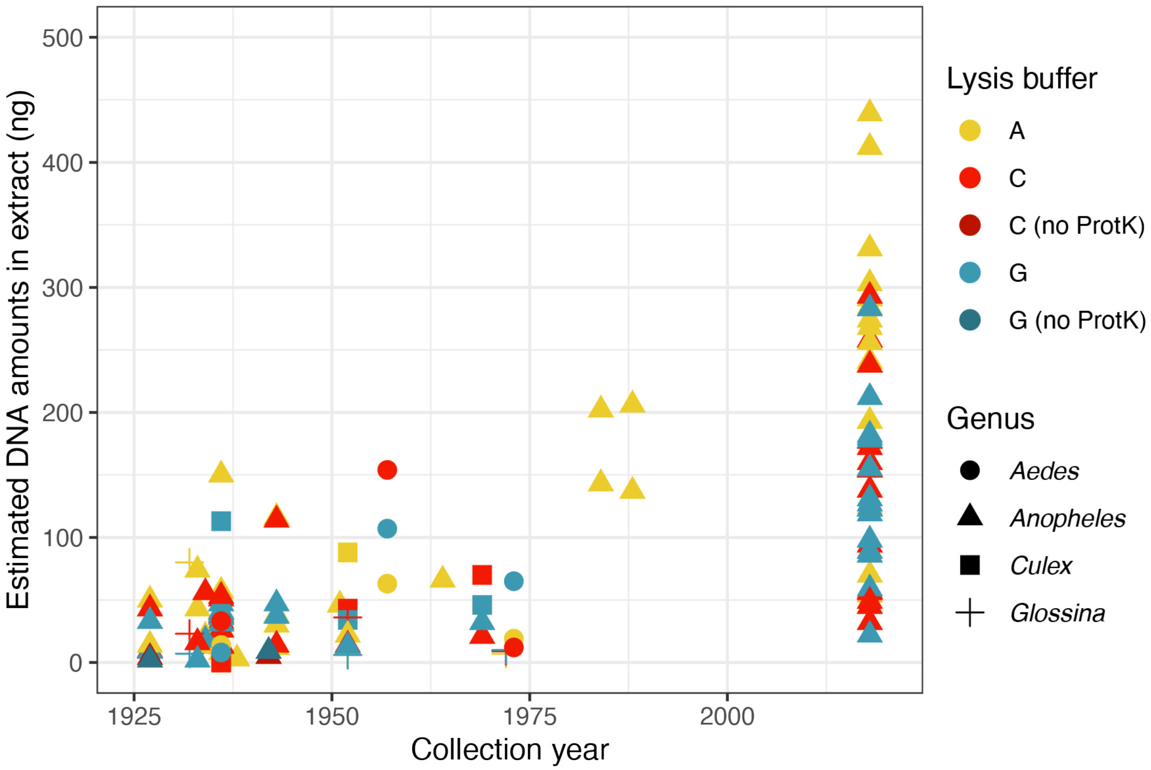


**Supplementary Figure S11.** Sample age effect on DNA yields. DNA amount estimates determined by Quant-iT™ PicoGreen™ dsDNA Assay Kit and ordered by their collection year for all historic Diptera processed in this study compared to estimates for dried present-day *Anopheles* specimens extracted using the same three DNA lysis buffers (A, C, G) and a MinElute 96 UF PCR Purification Kit (data from (Makunin et al. 2021)).

**NHMUK014063479 *Anopheles funestus*, overnight buffer A**

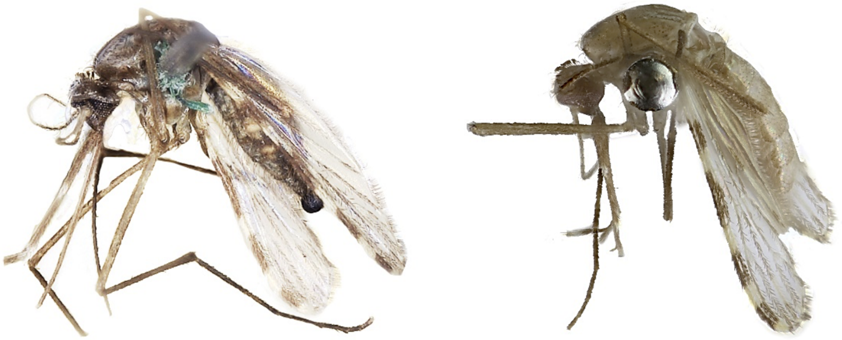

**NHMUK014063467 *Anopheles funestus*, overnight buffer C**

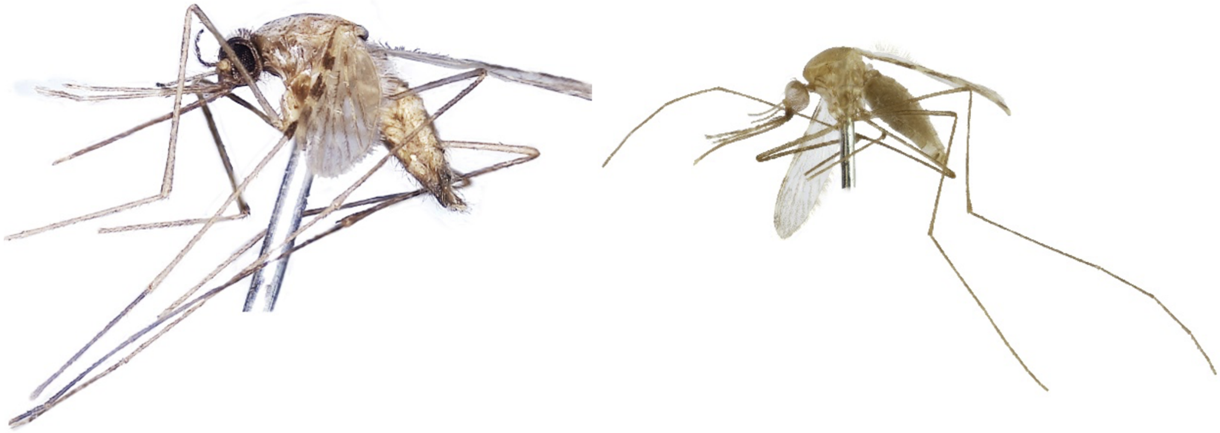

**NHMUK014063604 *Anopheles funestus*, overnight buffer G**

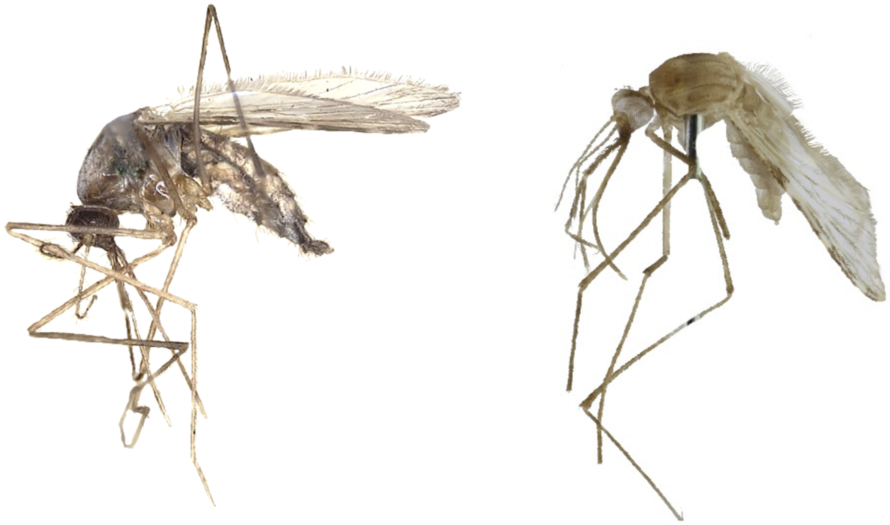

**Supplementary Figure S12.** Images of representative specimens before (left) and after (right) DNA extraction and CPD: *Anopheles funestus* overnight extractions with lysis buffers A (top), C (middle) and G (bottom). Images were processed using Adobe Photoshop to remove backgrounds and correct brightness/contrast.

**NHMUK013655441 *Anopheles funestus*, 2 h buffer A**

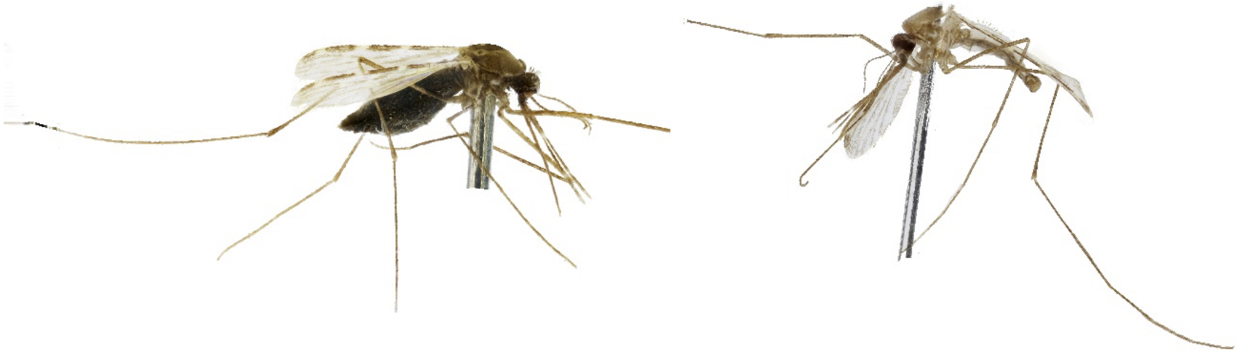

**NHMUK013655439 *Anopheles funestus*, 2 h buffer C**

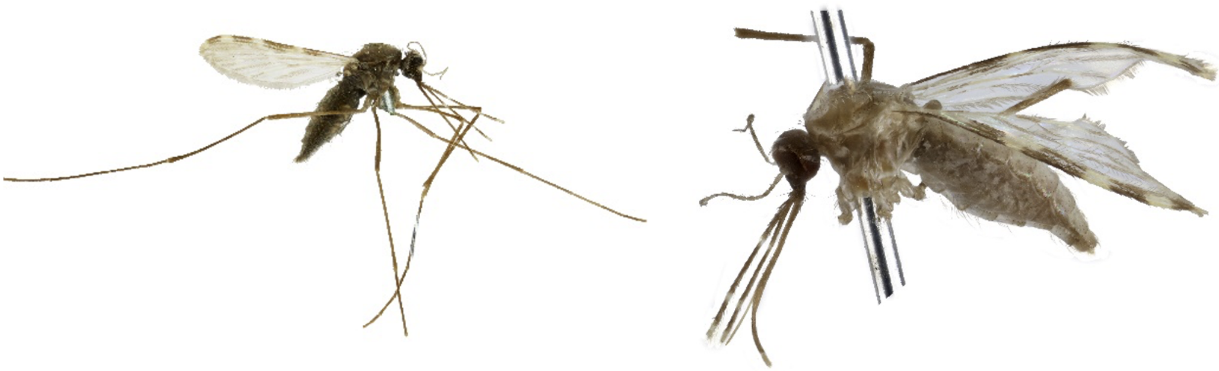

**NHMUK013655440 *Anopheles funestus** (actually *An. rivulorum*), 2 h buffer G**

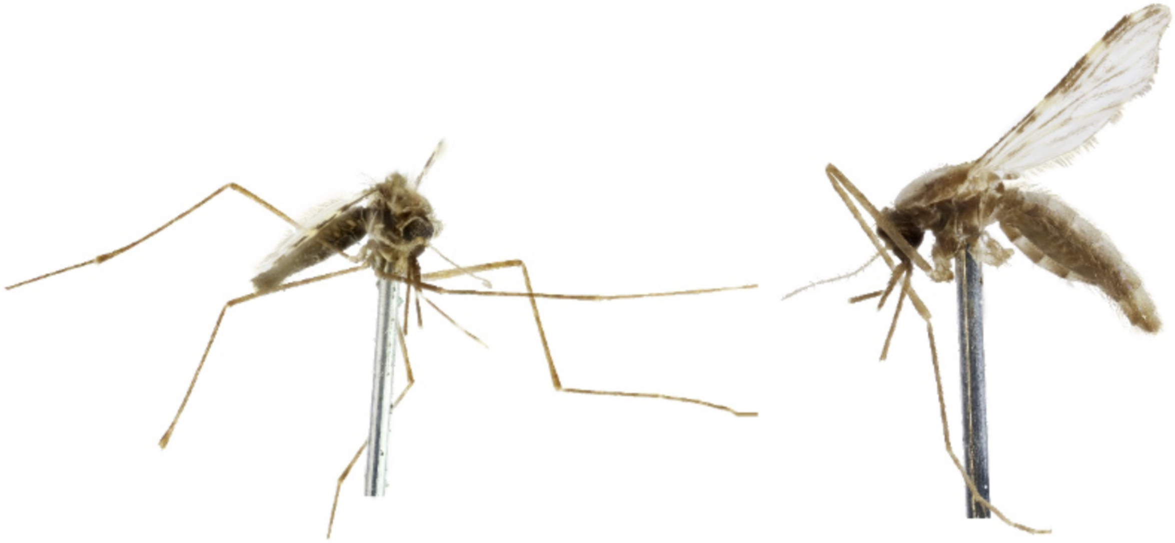

**Supplementary Figure S13.** Images of representative specimens before (left) and after (right) DNA extraction and CPD: *Anopheles funestus* 2 h extractions with lysis buffers A (top), C (middle) and G (bottom). Images were processed using Adobe Photoshop to remove backgrounds and correct brightness/contrast.

**NHMUK010629159 *Aedes aegypti*, 2 h buffer A, pass**

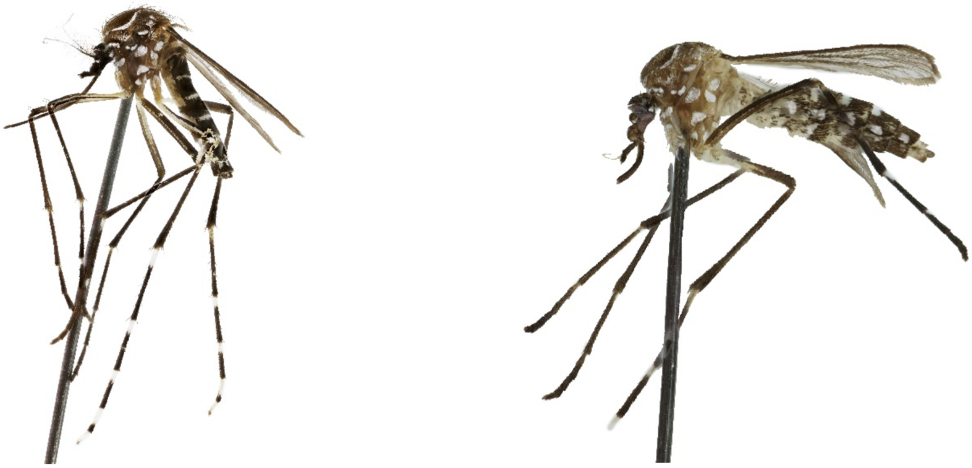

**NHMUK010629182 *Aedes aegypti*, 2 h buffer C, pass**

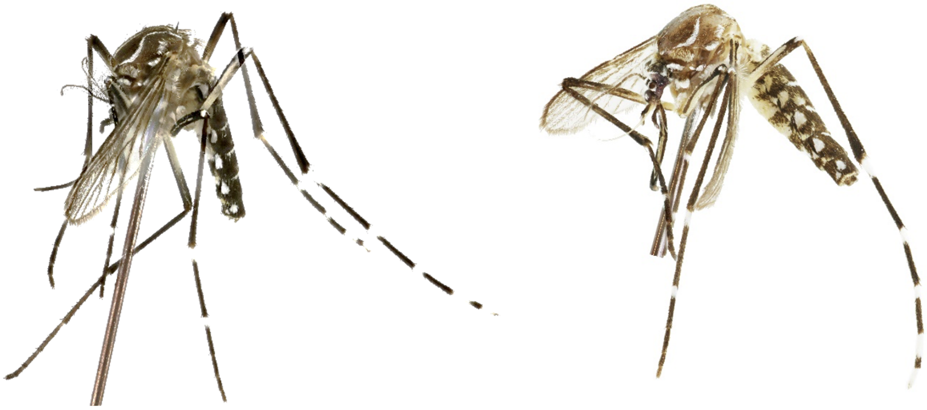

**NHMUK010629169 *Aedes aegypti*, 2 h buffer G, fail**

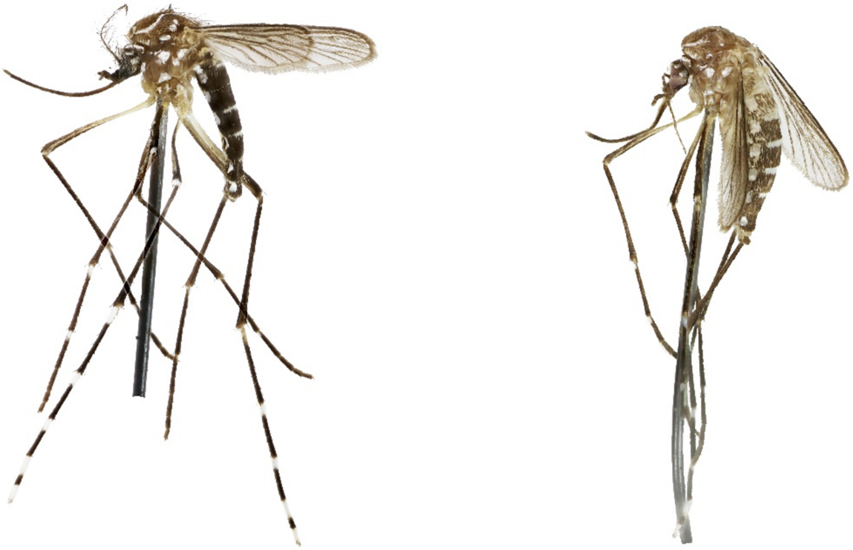

**Supplementary Figure S14.** Images of representative specimens before (left) and after (right) DNA extraction and CPD: *Aedes aegypti* 2 h extractions with lysis buffers A (top), C (middle) and G (bottom). Images were processed using Adobe Photoshop to remove backgrounds and correct brightness/contrast.

**NHMUK010633501 *Anopheles gambiae*, 2 h buffer A, fail**

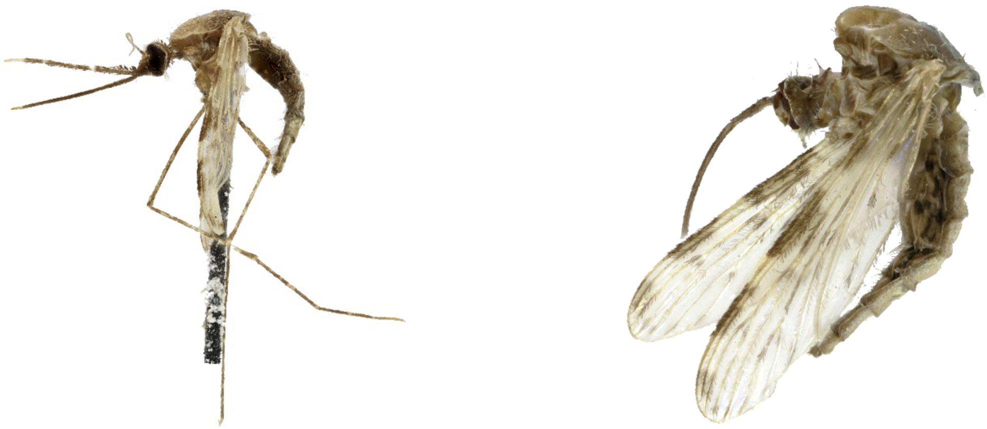

**NHMUK010633503 *Anopheles gambiae*, 2 h buffer G, fail**

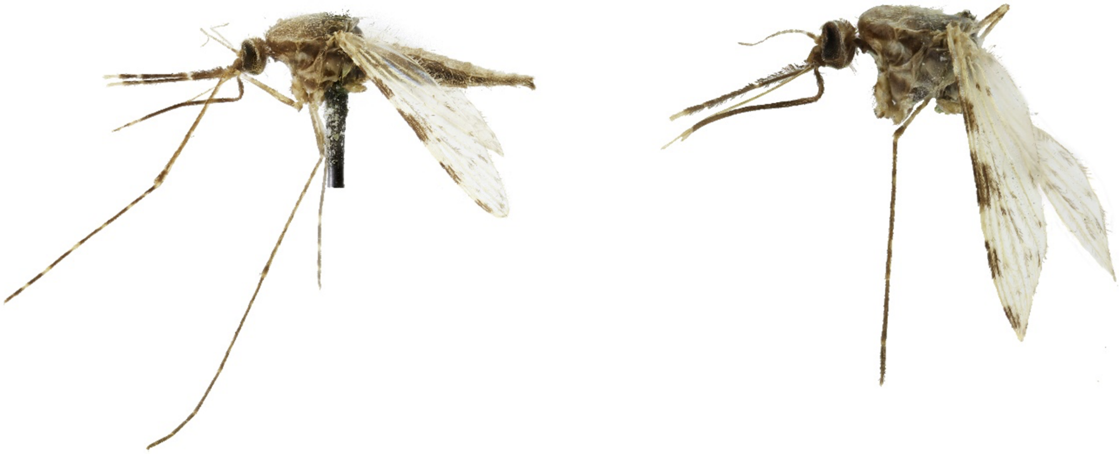

**Supplementary Figure S15.** Images of representative specimens before (left) and after (right) DNA extraction and CPD: *Anopheles gambiae* 2 h extractions with lysis buffers A (top), and G (bottom). Lysis buffer C is showcased in Figure 3A. Images were processed using Adobe Photoshop to remove backgrounds and correct brightness/contrast.

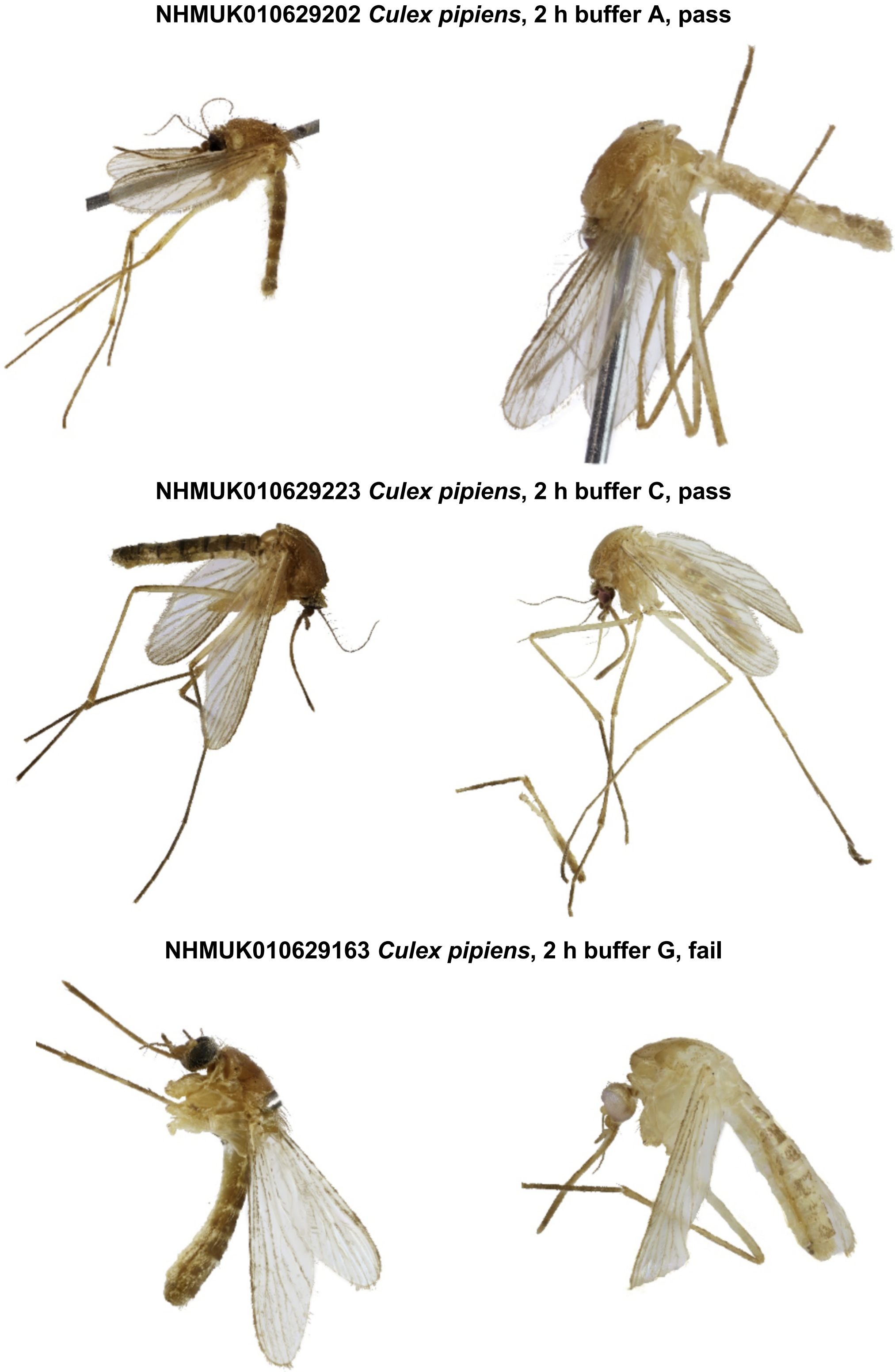


**Supplementary Figure S16.** Images of representative specimens before (left) and after (right) DNA extraction and CPD: *Culex pipiens* 2 h extractions with lysis buffers A (top), C (middle) and G (bottom). Images were processed using Adobe Photoshop to remove backgrounds and correct brightness/contrast.

**NHMUK010629183 *Glossina morsitans*, 2 h buffer A, pass**

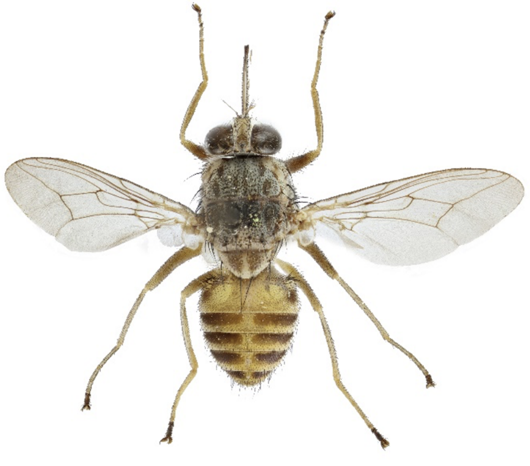

**NHMUK010629158 *Glossina morsitans*, 2 h buffer C, pass**

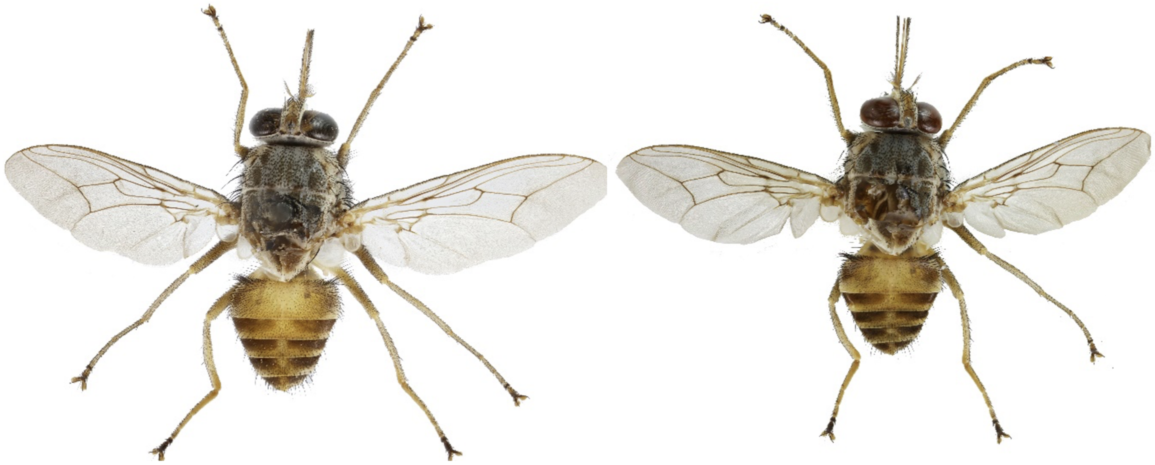

**NHMUK010629154 *Glossina morsitans*, 2 h buffer G, pass**

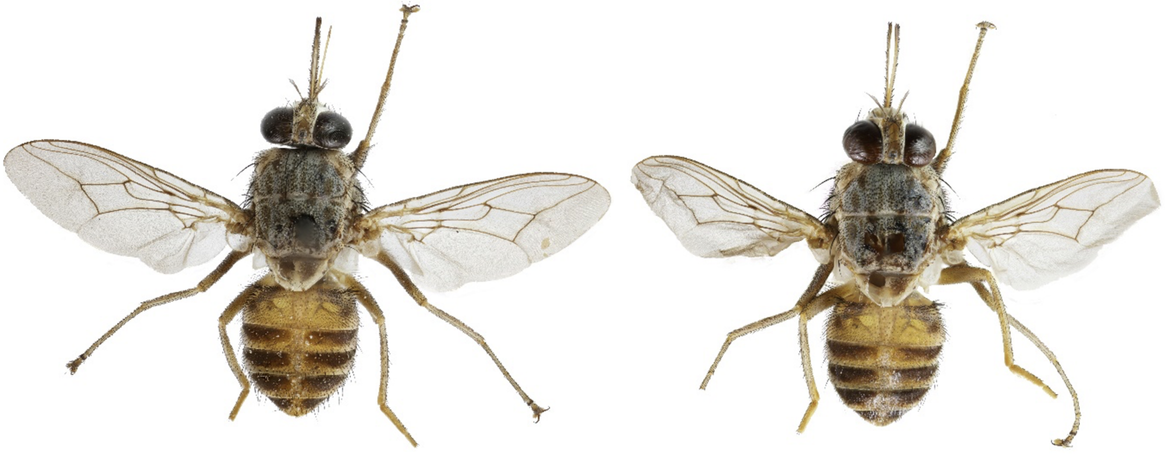

**Supplementary Figure S17.** Images of representative specimens before (left) and after (right) DNA extraction and CPD: *Glossina morsitans* 2 h extractions with lysis buffers A (top), C (middle) and G (bottom). Images were processed using Adobe Photoshop to remove backgrounds and correct brightness/contrast.

**Supplementary Table S1. (separate file)** Verbatim London Natural History Museum (NHM) labels associated with each specimen in this study.

**Supplementary Table S2. (separate file)** Assessment of morphological changes post-DNA extraction and CPD for *Aedes*, *Anopheles*, *Culex* and *Glossina* specimens from our final experiment.

**Supplementary Table S3. (separate file)** Overview of DNA extraction, library preparation and sequencing data (where relevant) for all specimens in this study.

**Supplementary Table S4.** Known voltage-gated sodium channel (VGSC) mutations that lead to insecticide resistance in present-day *An. gambiae/coluzzii/arabiensis* populations (Clarkson et al. 2021) evaluated in our historic *An. gambiae* and *An. arabiensis* specimens. For each position we note down the observed base and the number of sequences in brackets, with n.a. for positions where no sequences were present.

| **Position** | **Variant** | **NHMUK 010633470** | **NHMUK 010633474** | **NHMUK 010633475** | **NHMUK 010633479** | **NHMUK 010633480** | **NHMUK 010633481** | **NHMUK 010633482** | **NHMUK 012805003** | **NHMUK 012805007** | **NHMUK 012805014** | **NHMUK 012804999** | **NHMUK 012805000** | **NHMUK 012805015** |
|---|---|---|---|---|---|---|---|---|---|---|---|---|---|---|
| **2,390,177** | **G>A** | G(1) | G(1) | G(2) | G(2) | G(1) | n.a. | G(2) | G(2) | n.a. | G(1) | n.a. | G(7) | G(11) |
| **2,391,228** | **G>C,T** | n.a. | G(2) | G(1) | n.a. | G(2) | G(3) | G(1) | G(2) | G(1) | G(1) | G(2) | G(9) | G(13) |
| **2,399,997** | **G>C** | G(2) | G(2) | n.a. | G(3) | G(5) | G(1) | G(1) | n.a. | n.a. | G(3) | G(2) | G(17) | G(8) |
| **2,400,071** | **G>A,T** | G(3) | n.a. | n.a. | n.a. | G(2) | n.a. | G(3) | G(2) | G(1) | G(3) | n.a. | G(17) | G(7) |
| **2,402,466** | **G>T** | G(2) | G(1) | n.a. | G(1) | G(1) | n.a. | G(1) | n.a. | n.a. | G(2) | G(3) | G(11) | G(7) |
| **2,407,967** | **A>C** | A(3) | n.a. | A(1) | A(1) | n.a. | A(4) | A(3) | A(2) | A(1) | A(1) | A(2) | A(14) | A(14) |
| **2,416,980** | **C>T** | C(1) | n.a. | C(1) | C(1) | C(3) | C(1) | C(1) | C(1) | C(1) | C(5) | C(1) | C(15) | C(10) |
| **2,422,651** | **T>C** | T(1) | T(1) | T(1) | T(1) | T(2) | T(2) | T(2) | n.a. | n.a. | T(2) | n.a. | T(9) | T(9) |
| **2,422,652** | **A>T** | A(1) | A(1) | A(1) | A(1) | A(2) | A(2) | A(2) | n.a. | n.a. | A(2) | n.a. | A(10) | A(9) |
| **2,429,617** | **T>C** | T(3) | T(1) | T(1) | n.a. | n.a. | T(1) | n.a. | T(1) | n.a. | T(4) | T(1) | T(16) | T(10) |
| **2,429,745** | **A>T** | A(2) | A(2) | A(3) | n.a. | A(2) | n.a. | n.a. | A(4) | n.a. | A(1) | A(2) | A(12) | A(10) |
| **2,429,897** | **A>G** | A(1) | A(1) | n.a. | A(1) | n.a. | n.a. | A(4) | n.a. | n.a. | A(2) | A(1) | A(13) | A(16) |
| **2,429,915** | **A>C** | n.a. | A(1) | n.a. | A(1) | n.a. | A(1) | A(3) | n.a. | n.a. | A(2) | n.a. | A(14) | A(19) |
| **2,430,424** | **G>T** | G(1) | G(1) | n.a. | G(2) | G(3) | G(3) | G(2) | G(1) | G(1) | G(3) | G(2) | G(7) | G(14) |
| **2,430,817** | **G>A** | G(2) | n.a. | G(1) | G(5) | G(4) | G(3) | n.a. | G(1) | G(2) | G(1) | n.a. | G(16) | G(10) |
| **2,430,863** | **T>C** | T(1) | T(2) | T(3) | T(1) | T(1) | T(1) | T(1) | T(2) | n.a. | T(2) | T(2) | T(16) | T(9) |
| **2,430,880** | **C>T** | C(2) | C(2) | C(3) | C(1) | C(1) | C(2) | C(1) | C(1) | n.a. | C(3) | C(3) | C(13) | C(9) |
| **2,430,881** | **C>T** | C(2) | C(2) | C(3) | C(1) | C(1) | C(2) | C(1) | C(2) | n.a. | C(3) | C(3) | C(13) | C(10) |
| **2,431,019** | **T>C** | T(1) | T(1) | n.a. | n.a. | n.a. | T(1) | T(3) | T(5) | T(2) | T(4) | T(4) | T(14) | T(6) |
| **2,431,061** | **C>T** | C(1) | C(2) | n.a. | n.a. | C(3) | C(3) | C(3) | C(1) | n.a. | C(1) | C(5) | C(11) | C(9) |
| **2,431,079** | **T>C** | T(1) | T(2) | T(2) | n.a. | T(4) | T(4) | T(2) | T(3) | n.a. | T(2) | T(4) | T(10) | T(8) |
